# Neural representational geometries correlate with behavioral differences in monkeys and recurrent neural networks

**DOI:** 10.1101/2022.10.05.511024

**Authors:** Valeria Fascianelli, Aldo Battista, Fabio Stefanini, Satoshi Tsujimoto, Aldo Genovesio, Stefano Fusi

**Affiliations:** Center for Theoretical Neuroscience, Columbia University, NY, USA; Zuckerman Mind Brain Behavior Institute, Columbia University, NY, USA; Center for Neural Science, New York University, NY, USA; SixthFactor Pte. Ltd, Singapore; Department of Physiology and Pharmacology, Sapienza University of Rome, Rome, Italy; Department of Neuroscience, Vagelos College of Physicians and Surgeons, Columbia University Irving Medical Center, NY, USA; Kavli Institute for Brain Science, Columbia University, NY, USA

**Keywords:** representational geometry, abstraction, disentangled representations, individual differences, behavioral differences, strategy, dorsolateral prefrontal cortex, recurrent neural networks

## Abstract

Animals likely use a variety of strategies to solve laboratory tasks. Traditionally, combined analysis of behavioral and neural recording data across subjects employing different strategies may obscure important signals and give confusing results. Hence, it is essential to develop techniques that can infer strategy at the single-subject level. We analyzed an experiment in which two monkeys performed a visually cued rule-based task. The analysis of their performance shows no indication that they used a different strategy. However, when we examined the geometry of stimulus representations in the state space of the neural activities recorded in dorsolateral prefrontal cortex, we found striking differences between the two monkeys. Our purely neural results induced us to reanalyze the behavior. The new analysis showed that the differences in representational geometry correlate with differences in the reaction times, revealing behavioral differences we were unaware of. All these analyses indicate that the monkeys are using different strategies. Finally, using recurrent neural network models trained to perform the same task, we show that these strategies correlate with the amount of training, suggesting a possible explanation for the observed neural and behavioral differences.

## Introduction

Although the tasks designed in a laboratory are relatively simple and are performed in highly controlled situations, different animals can still adopt different strategies to solve the same task. It is surprisingly difficult to reproduce the exact same behavior in different laboratories, even when the training protocol, the experimental hardware, software, and procedures are standardized [1]. In many situations, it is also possible that the behavioral performance is the same, but the strategy used to perform the task is different. Consider, for example, a task in which multiple stimulus properties must be mapped onto appropriate behavioral responses. Such a task can be accomplished by rote learning of this map, but if the task involves structure across stimulus attributes, including irrelevant stimulus features, learning can be simplified by adopting more “intelligent” strategies that exploit this structure. All these strategies may produce the same level of task performance, so how can we reveal them?

Here, we show that this can be done by examining the geometry of stimulus representations in the state space of recorded neural activities in dorsolateral prefrontal cortex. The recorded neural responses are typically very diverse and seemingly disorganized [2, 3, 4, 5]. However, when the neural activity is analyzed at the population level, it is often possible to identify interesting and informative “structures”. In particular, the analysis of the geometry of the neural representations has recently revealed that some variables are represented in a special format that enables generalization to novel situations [6]. The representational geometry is defined by the distances between points representing different experimental conditions in the neural activity space. The set of points of all the experiment conditions defines an object with specific computational properties [7], which can be preserved across subjects [8]. For example, if the points define a high-dimensional object (in this article, we always consider the embedding dimensionality when we speak about dimensionality), then a linear decoder can separate the points in a large number of ways, permitting a downstream neuron to perform many different tasks [2, 3]. If, instead, the points define a low-dimensional object, the representations allow a simple linear decoder of one variable to generalize across the values of other variables [6]. These representations have been called abstract because of their generalization properties, and they are known as disentangled representations in the machine learning community [10, 11]. Abstract representations have been observed in several brain areas [6, 12, 13, 14, 15, 16, 17, 18, 19, 20, 21, 22], but it is still unclear whether they correlate with behavior. As pointed out by Krakauer et al. [23], a new conceptual framework that meaningfully maps the neural data to behavior is necessary to understand the brain-behavior relationship better, and to accomplish that, the analysis of the behavior should be as fine-grained as the analysis performed on the neural data.

Here we show that differences between neural representational geometries across subjects correlate with significant differences in their behavior, providing evidence that the aspects of the representational geometry we typically study could affect behavior. Thanks to these correlations, the analysis of the representational geometry is also an important tool for reliably interpreting individual differences in behavior.

More specifically, we analyzed the activity of neurons recorded in the dorsolateral prefrontal cortex (PFdl) of two monkeys performing a visually cued rule-based task [24]. The task required choosing between two spatial targets based on the rule cued by a visual stimulus, either staying with the same response as in the previous trial (after a stay cue) or shifting to the alternative response (after a shift cue). Crucially, the task average performance was the same for the two monkeys.

We systematically studied specific aspects of the geometry of neural representations. First, we looked at the ability of a linear decoder to classify all the task-relevant variables (the shape of the visual cue, the rule, the current and previous response). Then we tried to decode the variables that correspond to all the possible ways of dividing the conditions into two groups of equal size (balanced dichotomies). Some of these dichotomies correspond to obvious task-relevant variables, while others are not easily interpretable, but they may carry some meaning to the animal. We analyzed all the balanced dichotomies, and not only the four task-relevant variables, to assess which variables are decoded in an unbiased way, as introduced in Bernardi et al. [6]. We also computed the cross-condition generalization performance (CCGP) for all these dichotomies by training a decoder on a subset of conditions and testing on a different subset. These other conditions are completely novel for the decoder; hence, a high CCGP is possible only if the geometry allows for such generalization across conditions. Studying which variables have an elevated CCGP allowed us to identify which variables were represented in an abstract format and, therefore, describe another important aspect of the representational geometry [6]. This set of measures revealed that the representational geometry is strikingly different for the two monkeys: the first monkey is more “visual”, with a high decoding accuracy and high CCGP for the shape of the visual cues, whereas the second monkey is more “cognitive,” with a high decoding accuracy and high CCGP for the rule. This finding brought us to reanalyze the behavior, and we discovered that the reaction time patterns actually reflect the different representational geometries.

We then used Recurrent Neural Network (RNN) models to provide a possible mechanistic explanation for the origin of the differences in representational geometries and behaviors between the two monkeys. We trained multiple RNNs to perform the same cued rule-based task used in the experiment. Each RNN was randomly initialized and trained on a different sequence of stimuli. The training was interrupted when the RNN reached a certain level of performance. Although all the different RNNs performed equally well, different networks exhibited different representational geometries. The geometries of the networks that reached high performance earlier were similar to the geometry observed in the first monkey. The geometry of the RNNs that learned more slowly, resembled the one of the second monkey. Interestingly, when we then compared the reaction times of the RNNs and the monkeys, we observed the same patterns.

Our study demonstrates that the analysis of representational geometry enables us to discern individual differences in task-performing strategies. Furthermore, the compelling connection we established between the representational geometry and observed behavior underscores the critical role these geometric aspects might play in the execution of the task.

## Results

We analyzed single-unit recordings in the dorsolateral prefrontal cortex (PFdl) of two male rhesus monkeys. As the main message of this work is that the representational geometry can explain the differences in the behavior of the two monkeys, we will present the neural and behavioral results for each monkey separately. We refer to them as Monkey 1 and Monkey 2.

Both monkeys were trained to perform a visually cued rule-based task (Figure 1A). The task was to choose one of two spatial targets with a saccadic movement, according to the rule instructed in each trial by one of four possible visual cues (Figure 1B). Two cues instructed the monkey to “stay” with the target chosen in the previous trial, while the other two cues instructed to “shift” to the alternative target. In each trial, the visual cue was randomly chosen. At the time of the recordings, both monkeys were already trained, and they were performing the task with the same high accuracy.

**Fig. 1:**
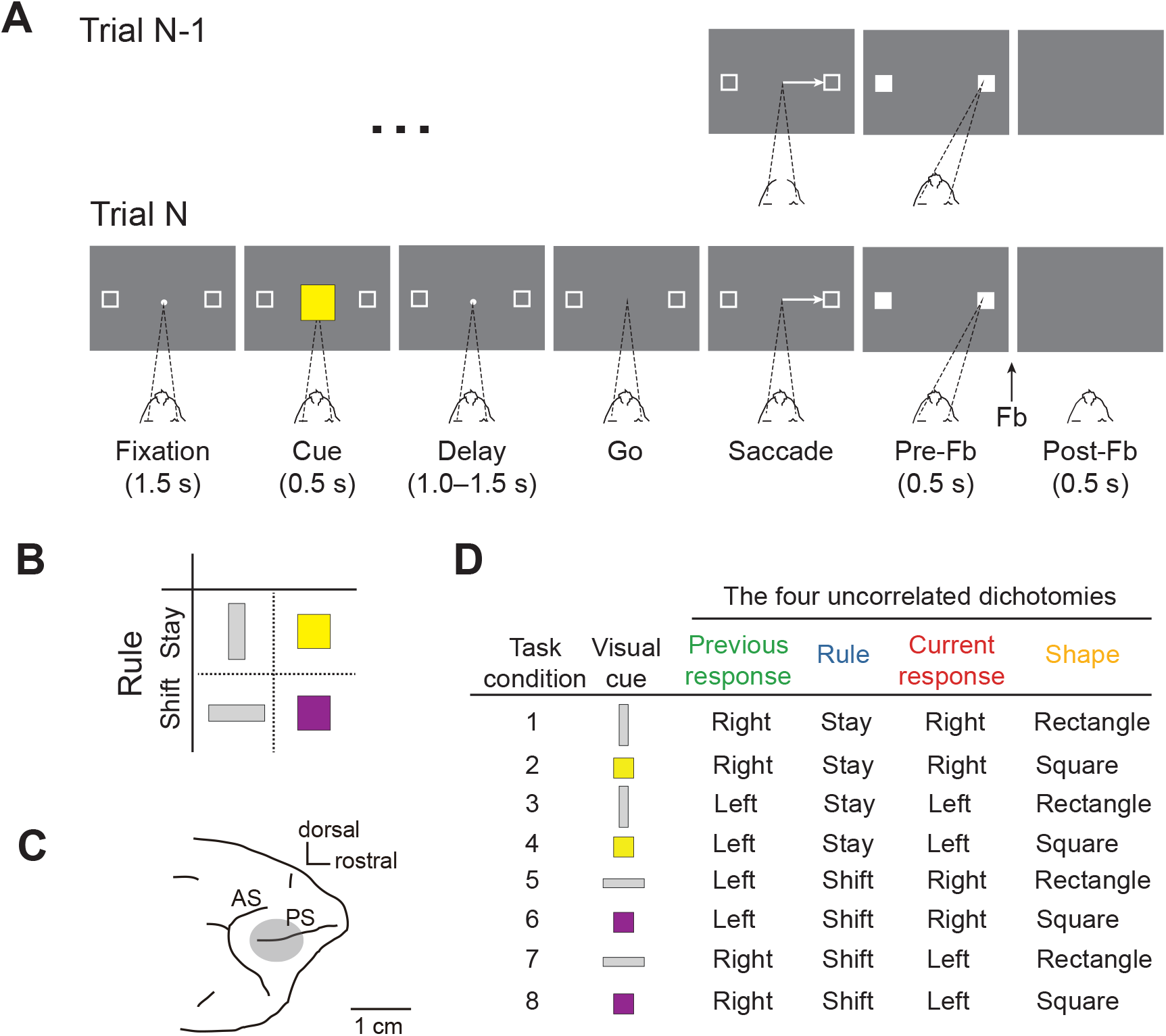
Behavioral task, visual cues, recording site, and task conditions. **A)** Example of two consecutive trials of the visually cued rule-based task with temporal ordering of task events from left to right. The dark gray rectangle represents the video screen as viewed by the monkey. The target of the monkey’s gaze is indicated by dashed lines. In this example, trial N is a stay trial instructed by the yellow square, requiring the monkey to choose the same target (right) chosen in the previous trial N-1. Fb, Feedback. **B)** Visual cues presented to the monkey. Each visual cue instructed the rule to be applied: the vertical grey rectangle and yellow square instructed the stay rule; the horizontal grey rectangle and purple square instructed the shift rule. **C)** Recording area in dorsolateral prefrontal cortex. AS, Arcuate Sulcus; PS, Principal Sulcus. **D)** List of the eight task conditions defined as the combination of the four main uncorrelated dichotomies: previous response (green), rule (blue), current response (red), and shape of the visual cue (orange). The color code of the four dichotomies is conserved across all the figures.

We found significant differences between the two monkeys when we analyzed the geometry of the neural representations recorded during the task. The representational geometry is defined by the set of distances between the points in the firing rate space that represent different conditions (see, e.g., [25]). This is a relatively large set of variables, which are not defined in a unique way as there are several reasonable measures of distances in the presence of noise. We found significant differences between the two monkeys by focusing on two particular aspects of the geometry that also have the advantage of being crossvalidated and interpretable: the first is the set of linear decoding accuracies for the task-relevant variables and all the other variables that correspond to balanced dichotomies of the conditions (i.e., all the possible ways of dividing the conditions into two equal groups). The task-relevant variables are the previous response, the rule, the current response, and the shape of the visual cue (Figure 1D). The latter identifies whether the visual cue is a rectangle or a square, although the cue also differs because the rectangles are grey and the squares are colored (yellow and purple).

The second aspect of the geometry is related to the ability of a linear classifier to generalize across conditions when trained to decode the balanced dichotomies (cross-condition generalization performance or CCGP [6]).

The decoding accuracy is directly related to the distance between two groups of points, and in this respect, it is a geometrical measure. It is better than the average distance because it is cross-validated and it takes into account the structure of the noise, similar to the Mahalanobis distance [26]. Moreover, it is interpretable because it tells us something about the represented variables. The second quantity, the CCGP, is more sensitive to the angles between coding directions, another aspect of the representational geometry: the ability of a linear classifier to generalize depends on the parallelism of the coding directions [6]. Say we consider two binary variables *x* and *y*, and we train a decoder to report the value of variable *x* from the patterns of neural activity. If this decoder is trained only in the situations in which *y* = *y*_1_, it is not guaranteed to work right away for a different value of *y*, say *y* = *y*_2_. In order to generalize to *y* = *y*_2_, it is necessary that the coding direction of *x* (i.e., the direction from the points corresponding to neural activities when *x* = *x*_1_ to *x* = *x*_2_) for *y* = *y*_1_ is approximately the same as for *y* = *y*_2_. CCGP also takes into account the noise structure, and it is crossvalidated. Moreover, if a variable has high CCGP it means that the variable is encoded in a special format that we formerly defined as “abstract”[6]. The variable is encoded in an abstract format (or simply, it is abstract) because the coding direction does not depend on the specific instance. This guarantees special generalization properties (cross-condition), which are the hallmark of abstraction.

### Differences between the neural representational geometries of the two monkeys

Our neural database consists of 289 and 262 neurons from Monkey 1 and Monkey 2, respectively. To investigate which task variables can be decoded, we built pseudo-simultaneous trials (pseudo trials) for each monkey separately (see Methods). We defined the pseudo trial as the combination of spike counts randomly sampled from different trials of the same task condition [2]. For each neuron, the spike count was estimated in a 200*ms* time bin. We considered only neurons recorded for at least 5 complete and correct trials in each task condition for a total of 205/289 (71%) neurons for Monkey 1 and 188/262 (72%) neurons for Monkey 2.

We found that, in Monkey 1, almost all the dichotomies can be linearly decoded during the cue presentation (Figure 2A), but not all of them are in an abstract format, i.e., with a high CCGP (Figure 2B). Shape is the variable with the highest CCGP, followed in time by the current response, while the previous response and the rule can be decoded but do have a CCGP at chance, and hence are not in an abstract format. It is worth noting that the rule is represented in an abstract format in a later period, after the cue offset (Figure 2B). In Monkey 2, almost all the dichotomies can be linearly decoded during the cue presentation, except for the shape and the previous response (Figure 2C). The CCGP analysis reveals that, in Monkey 2, the rule is in an abstract format with the highest CCGP during the cue presentation, differently from Monkey 1 (Figure 2D). In both monkeys, instead, during the cue presentation, the current response is in an abstract format, while the previous response is not.

**Fig. 2:**
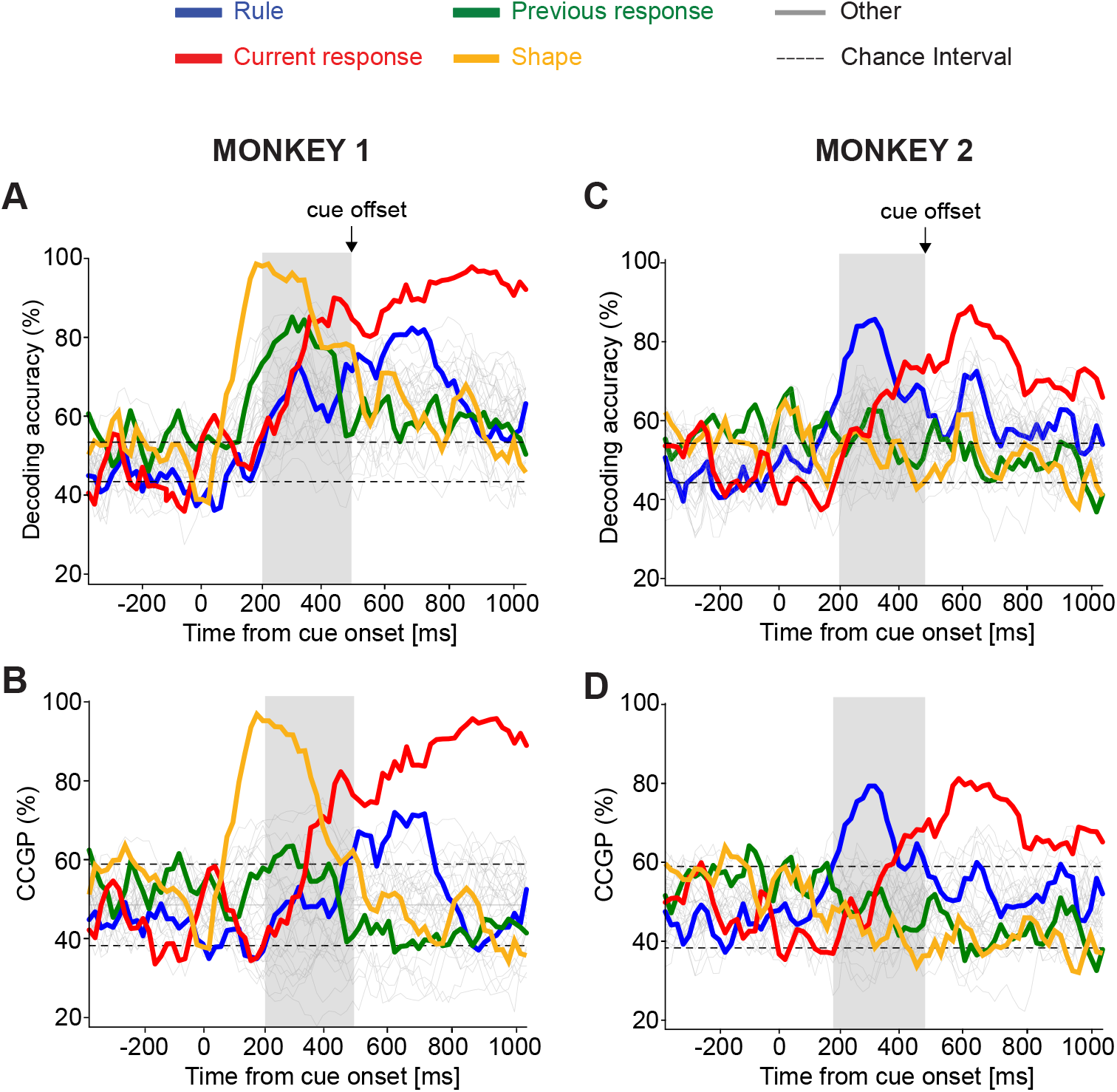
Decoding accuracy and CCGP as a function of time. Time is aligned to the cue onset lasting for 500*ms* (until the time of the cue offset indicated by the vertical black arrow). The horizontal dashed lines are *±*2 standard deviations of 100 cross-validations distribution obtained from null models. The grey vertical shade indicates the time bin starting at 200*ms* after cue onset until cue offset, in which we found the maximal difference between the neural representations of the two monkeys. **A)** Decoding accuracy of all the possible 35 dichotomies (i.e., all variables that correspond to grouping the conditions into two equal-size groups) in Monkey 1. During the cue presentation, most of the dichotomies can be decoded, in particular all the main task variables indicated with different colors. The shape of the visual cue (orange) is decoded with the highest accuracy, followed in time by the previous response (green), the current response (red), and the rule (blue). **B)** CCGP of the 35 dichotomies in Monkey 1. During the cue presentation, the shape (orange) is in an abstract format with the highest CCGP, followed in time by the current response (red). The Rule (blue) is not abstract during the cue presentation, but it becomes abstract after the cue offset. The previous response (green) is not in an abstract format. **C)** Decoding accuracy of all dichotomies in Monkey 2. During the cue presentation, the rule (blue) and the current response (red) can be significantly decoded. **D)** CCGP of all dichotomies in Monkey 2. Differently from Monkey 1, the rule (blue) is in an abstract format with the highest CCGP during cue presentation, followed in time by the current response (red). The shape (orange) and previous response (green) are not in an abstract format.

We performed an additional analysis to control whether the main differences in decoding accuracies between monkeys could depend on differences in the recording sites. Supplementary Figure 1A shows the location of the penetrations of the recordings in PFdl for both monkeys. It reveals an overlap between the recording sites in the two monkeys, particularly in the dorsal region to the principal sulcus, where all neurons in Monkey 2 were recorded. Thus, we examined the dorsal (106 neurons) and ventral (99 neurons) recordings separately for Monkey 1. When comparing the decoding accuracy of the task variables between neurons in the dorsal sites only (which match exactly with the sites in Monkey 2) with the combined dorsal and ventral recordings (see Figure 2A and Supplementary Figure 1B), we found comparable results. For this reason, we combined the recording sites in Monkey 1 in all the following analyses. The main difference between the dorsal and ventral recordings regards the representation of the previous response, which is encoded mainly in neurons in the ventral sites. This matches the previous response signal in Monkey 2, which is weakly decoded during the fixation period. It might be possible that even Monkey 2 could have a stronger previous response representation ventrally to the principal sulcus, but we lack the ventral recordings.

To better highlight the differences in the representational geometry, we focused our analysis on the 300*ms* time window in which the differences are large (from 200*ms* after the cue onset until the cue offset, grey vertical shade in Figure 2). The beeswarm plots in Figure 3A show the decoding accuracy and CCGP for all the possible dichotomies in the 300*ms* time window for Monkey 1 and Monkey 2. It is evident that Monkey 1 represents the shape of the visual cue in an abstract format (highest CCGP), while the rule is not abstract (CCGP at chance), even though it can be decoded (Figure 3A). The rule becomes abstract only later in the delay period after the cue offset (Figure 2B). Instead, for Monkey 2, the rule is the variable with the highest CCGP, while the shape of the visual cue is not abstract (CCGP at chance) (Figure 3A). Moreover, both monkeys represent the current response in an abstract format but not the previous response. Interestingly, in both monkeys, the current response is not abstract from the time when it can be decoded, but only slightly later (see Figure 2). These results suggest that Monkey 1 is grouping together the cues with the same shape, and hence it might be using a strategy based on the identity of individual visual stimuli. Instead, Monkey 2 might be using a more “cognitive” strategy because the rule is the variable with the highest decoding accuracy and CCGP, and hence Monkey 2 is grouping together the visual cues that correspond to the same rule, even though they are visually very different. We also assessed the dimensionality of the representation using the shattering dimensionality measure. The shattering dimensionality (SD) is defined as the average linear decoding performance for all possible balanced dichotomies. A high shattering dimensionality means that the linear decoder can perform a large number of inputoutput functions [2, 6]. We observed a higher value of SD in Monkey 1 than in Monkey 2 during the 300*ms* time window in which the differences in representational geometries are large (Figure 3A). A higher SD in Monkey 1 might be compatible with implementing a more “visual” strategy based on a lookup table, for which each visual cue is uniquely associated with a mapping from the previous response to the current response.

**Fig. 3:**
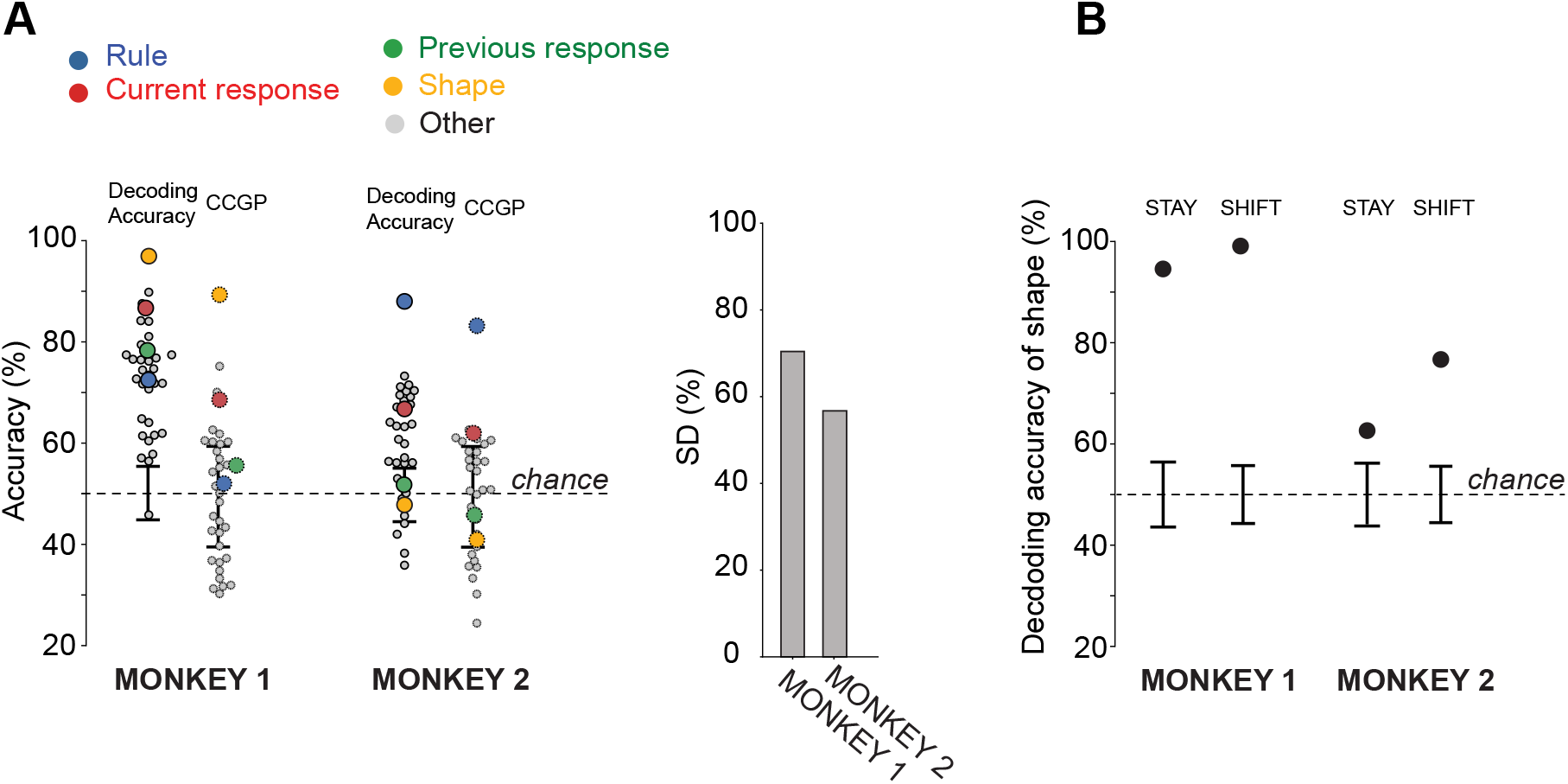
Summarizing the features of the representational geometry: decoding accuracy, CCGP, and shattering dimensionality (SD) during the last 300*ms* of cue presentation. **A)** Left: decoding accuracy (continuous-edge circles) and CCGP (dashed-edge circles) for each of the 35 dichotomies in the 300*ms* time bin during cue presentation in Monkey 1 and Monkey 2. Each circle is a different dichotomy. The four main dichotomies corresponding to task variables are highlighted with different colors. All the other dichotomies are in grey. Black error bars are the ±2 standard deviations around the chance level obtained from null models. Right: SD for Monkey 1 and Monkey 2 in the 300*ms* time bin during the cue presentation. The dimensionality of neural representation is higher in Monkey 1 than in Monkey 2. **B)** Decoding accuracy of shape in Monkey 1 and Monkey 2 in the stay and shift rule. The linear decoder was trained after projecting the neural activity of each pseudo trial in a lower dimensional space using a Multi-Dimensional Scaling (MDS) transformation. The shape is decodable in both rule conditions in both monkeys, though the accuracy is lower in Monkey 2.

Since the shape cannot be decoded in Monkey 2 using a linear classifier, we were wondering whether it is not encoded at all or if it could be decoded using other decoders. We decided to consider pairs of conditions separately, which is equivalent to considering non-linear decoders for all the points. Indeed, if two conditions are sufficiently separated, i.e., the distance between the corresponding points is large enough compared to the noise, then a linear decoder should work. This is true even when the dichotomy is not linearly separable. For example, in the case of XOR for four points that define a low-dimensional object like a square, a linear decoder would not be able to separate the two points on the diagonal from the other two, but it would separate all pairs of points if taken one pair at the time. In addition to considering pairs of points, we denoised the data by projecting the neural activity of a single pseudo trial into a lower dimensional space (3D) using the Multi-Dimensional Scaling (MDS) technique described in the Methods. Using this procedure, we found that the shape can be decoded in both monkeys. In particular, in Monkey 1, the shape can be decoded for both rule conditions with high accuracy (Figure 3B). This was expected as the shape was already linearly decodable for all the points without denoising (decoding accuracy in Figure 3A, left). Shape could also be decoded in Monkey 2, in both rule conditions (Figure 3B). These results show that both monkeys’ PFdl neurons encode the shape of the visual stimulus but with different geometries, making the shape linearly separable and in an abstract format only in one of the monkeys.

To visualize the different geometries of the two monkeys, we used the MDS transformation to reduce the dimensionality of the original representations. More specifically, we used MDS on the dissimilarity matrix containing the Euclidean distances between the average activity of two task conditions normalized by the variance along the direction that goes from one condition to the other (see Methods). Each point in the MDS plots is the average firing rate of each task condition in a 300*ms* time window during the cue presentation (Figure 4). For each monkey, we highlighted the different dichotomies (groups of conditions) by drawing lines between the conditions that are in the same group. In particular, the shape of the visual cue and current response are in an abstract format in Monkey 1, while the rule and current response are abstract in Monkey 2. For both monkeys, the current response is in an abstract format, while none have the previous response in an abstract format.

**Fig. 4:**
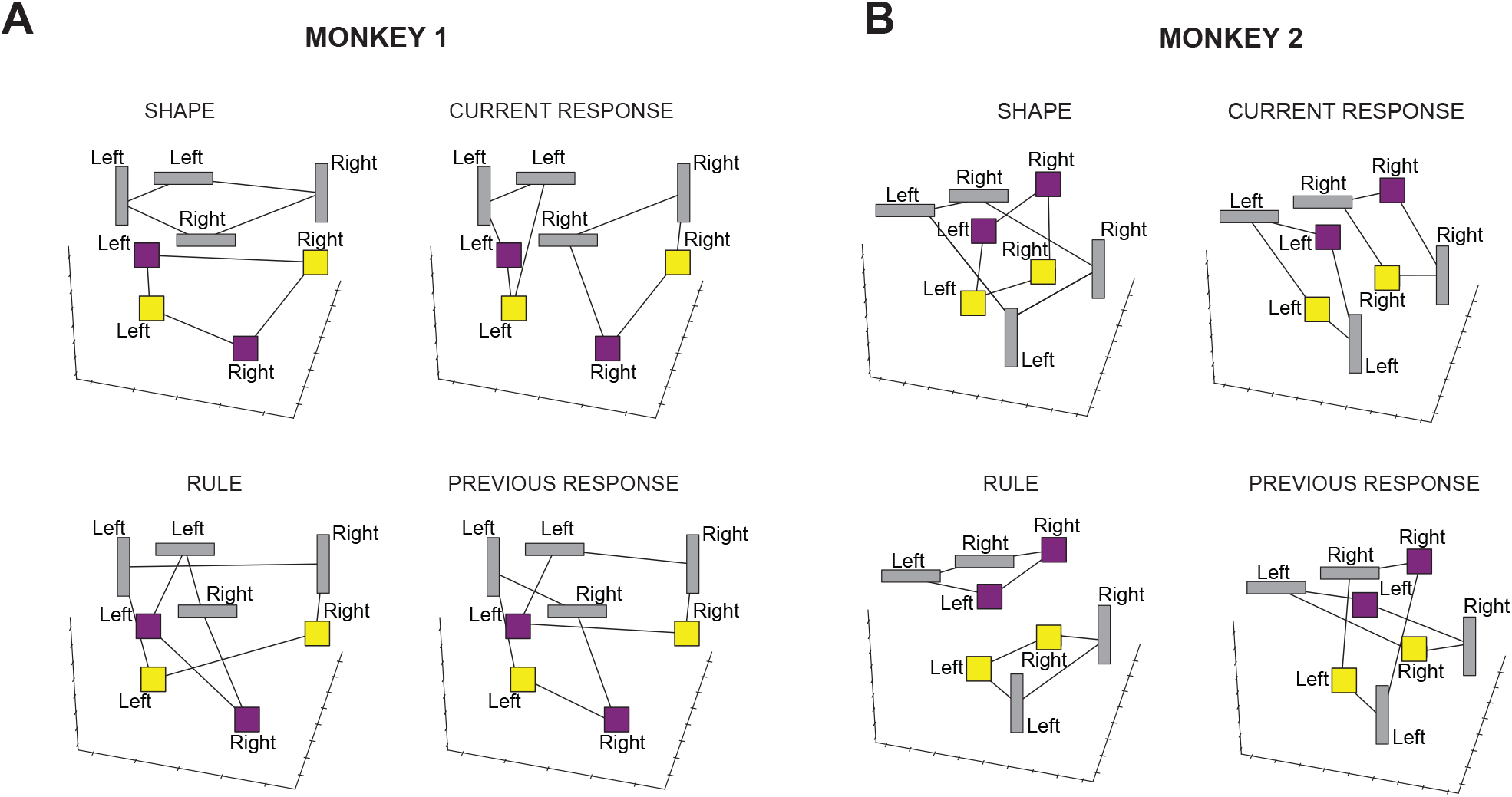
3-D Multi-Dimensional Scaling (MDS) plots. Each point represents the average firing rate in the 300*ms* time window during the cue presentation for one of the eight task conditions. The visual cue is shown along with the current response. The connecting black lines highlight the dichotomy indicated in the label above each plot. Shape and rule are the two dichotomies that mostly characterize the difference in representational geometry between the two monkeys, while the current response is in an abstract format for both monkeys. The previous response is not abstract in either of the monkeys. **A**) 3-D MDS plots in Monkey 1 for shape (top-left), current response (top-right), rule (bottom-left), and previous response (bottom-right). **B**) The same as in A) for Monkey 2.

### Behavioral differences between monkeys reflect differences in the neural representational geometry

The differences in the representational geometry are so striking that they induced us to reanalyze the behavior to look for more subtle individual differences. We analyzed 65 and 77 sessions for Monkey 1 and Monkey 2, respectively. As mentioned, we did not find any significant difference in the overall behavioral performance between the two monkeys (chi-square test, *p*-value=0.93; Figure 5A left). However, a significant difference emerged in the average reaction times (Mann-Whitney U test, *p*value=10^−15^; Figure 5A right). We then decided to analyze the behavior with finer-grained analyses. Indeed, since the neural analysis revealed a difference in the representational geometry of the shape and rule between the monkeys, we computed the average behavioral performance for each condition separately, by grouping the correct trials according to shape (rectangle and square) and rule (stay and shift). There is not a significant difference in the behavioral performance between different shapes (chi-square test: *p*-value=0.78 in Monkey 1, Figure 5B left; *p*-value=0.06 in Monkey 2, Figure 5D left), and between different rules (chi-square test: *p*-value=0.11 in Monkey 1, Figure 5B right; *p*-value=0.22 in Monkey 2, Figure 5D right) in both monkeys. Nevertheless, a significant difference in reaction times emerged across conditions in each monkey. In particular, Monkey 1, with high decoding accuracy and CCGP for the shape, has an average reaction time that significantly changes with the shape of the visual cue (Mann-Whitney U test: *p*-value = 0.002; Figure 5C, left) regardless of the rule (Mann-Whitney U test: *p*-value = 0.05; Figure 5C, right). On the opposite, Monkey 2, with high decoding accuracy and CCGP for the rule, shows an average reaction time that significantly changes with the rule (Mann-Whitney U test: *p*-value = 10^−10^; Figure 5E right) regardless of the shape (Mann-Whitney U test: *p*-value = 0.28; Figure 5E left).

**Fig. 5:**
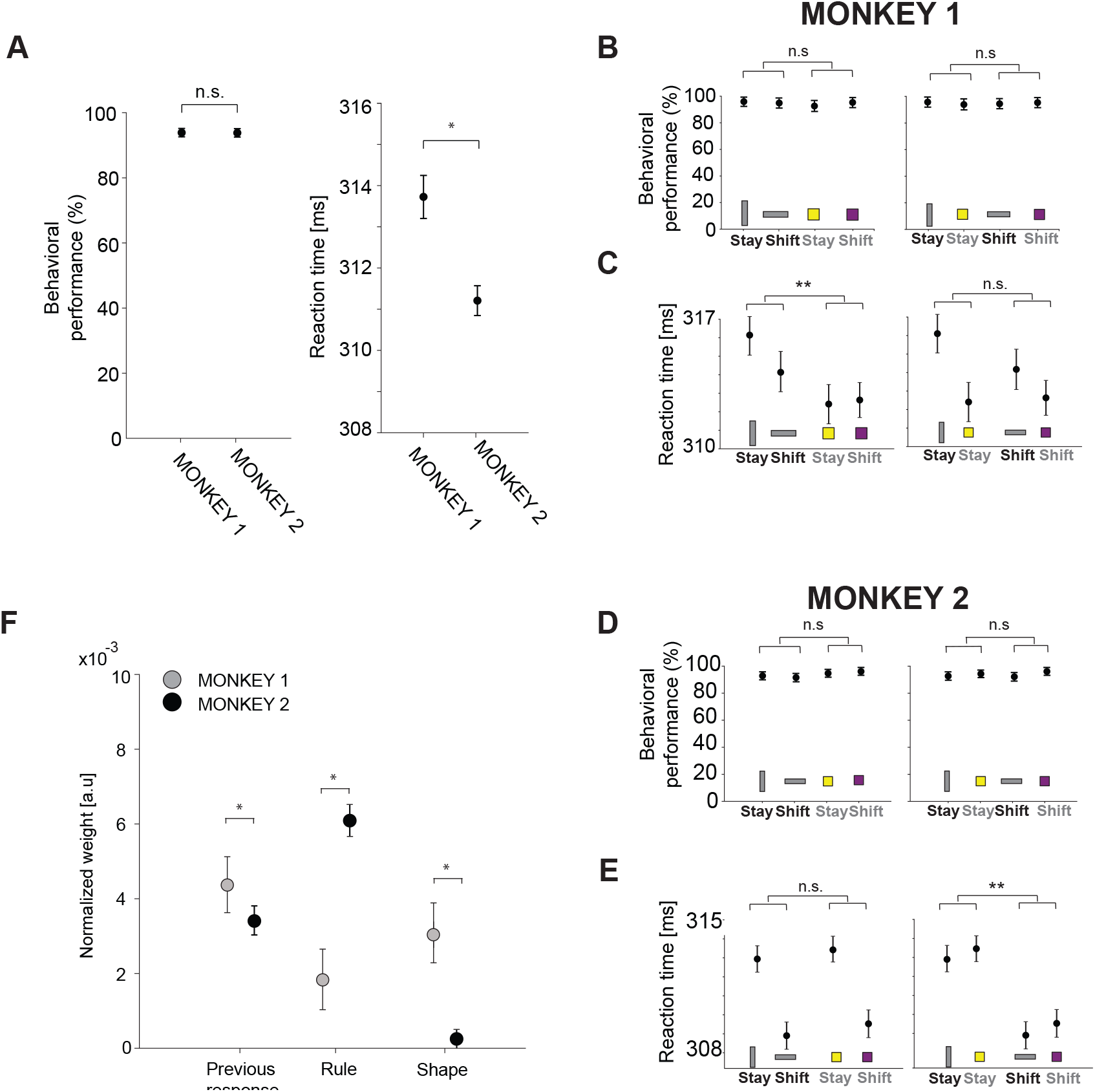
Behavioral performance, reaction times of the monkeys, and multi-linear regression behavioral model reflecting differences in the neural representational geometries. **A)** Left: average behavioral performance across sessions for Monkey 1 and Monkey 2. Both monkeys performed the task with high accuracy. The error bars indicate the confidence interval at 95% of confidence level. Right: average reaction time across sessions for Monkey 1 and Monkey 2. A significant difference emerged in the average reaction time between the two monkeys. The error bars are the standard error of the mean. n.s. not significant: chi-square test, *p*-value*>*0.05. *: Mann-Whitney U test, *p*-value*<*0.05. **B)** Mean behavioral performance across sessions for Monkey 1 computed separately for each rule and shape. The x-axis indicates the rule, and the y-axis is the mean performance averaged across sessions. The visual cue of each condition is indicated at the bottom of the plot. On the left(right), the visual cue order reflects shape(rule). n.s. not significant: chi-square test, *p*-value*>*0.05. **C)** Mean reaction time across sessions for Monkey 1. As in B), the x-axis indicates the rule, and the y-axis is the reaction time averaged across sessions. The error bar is the standard error of the mean. n.s. not significant: Mann-Whitney U test, *p*-value*>*0.05; **: Mann-Whitney U test, *p*-value*<*0.01. **D)** The same as in B) but for Monkey 2. **E)** The same as in C) but for Monkey 2. **F)**Weights of the three independent factors predicting the reaction time of single trial in a multi-linear regression model. Weights are normalized to the maximum weight that is the previous response and rule interaction term in both monkeys (see Supplementary Figure 2). The error bars are the 2 standard deviations of weights across of 100 models. The variance explained (r-squared) by the models is 12% and 18% for Monkey 1 and Monkey 2, respectively. *: Mann-Whitney U test, p*<*0.05.

The differences in reaction times are significant and they nicely reflect the representational geometry, but they are relatively small. So we decided to further investigate the behavior to see whether these differences could be predicted by looking at the recent series of events and monkey responses. In particular, we fitted a multi-linear regression model to predict the reaction time on a trialby-trial basis using three factors: the previous response, the shape of the visual cue, and the rule. We also considered all the interaction terms (see Supplementary Figure 2). We found that the rule factor has a stronger weight in predicting reaction times in Monkey 2 than in Monkey 1 (Mann-Whitney U test: *p*-value=10^−34^; Figure 5F). Vice versa, the shape is a stronger factor in predicting the reaction time of Monkey 1 (Mann-Whitney U test: *p*-value=10^−34^; Figure 5F). Supplementary 2 shows that the strongest factor in predicting the reaction time is the interaction of the previous response and the rule in both monkeys because the combination of these two factors is essential for choosing the correct response.

### Correlation between representational geometries and reaction times in recurrent neural networks

To investigate the origin of the differences in representational geometries between the two monkeys and the correlation with reaction times, we trained vanilla Recurrent Neural Networks (RNNs) to perform the visually cued rule-based task through deep reinforcement learning algorithms [27]. Artificial neural networks have been shown to be a powerful tool for understanding the normative aspects of neural representations [28], especially when trained with reinforcement learning, which resembles the protocols used to train animals [29, 30]. As these networks are often over-parametrized, numerous choices of parameters correspond to the same task performance. When these networks are trained multiple times, the solutions found by the same training algorithm can differ substantially, leading to networks that might even implement qualitatively different strategies to solve the same problem. This is usually considered an inconvenience, as much as the individual differences observed across animals. In our case, we took advantage of this variability across simulations to understand the relation between different geometries and the reaction times that were observed in the experiments.

We trained 80 RNNs using the Proximal-Policy-Optimization (PPO) as a deep reinforcement learning algorithm [27]. The architecture of the network, the learning algorithm, and the statistics of the inputs were the same for all the RNNs. However, the networks’ weights and biases were initialized randomly, using a different realization for every network. The inputs were: 1) the visual cue, encoded by two one-hot vectors of three units each, with the first vector encoding the shape, and the second vector the color; 2) the previous response, encoded by one one-hot vector of two units; 3) the fixation input, a scalar that instructs the network either to maintain fixation or not (Figure 6A). The inputs were passed through fixed, non-trained, random weights to an expansion layer of 100 Rectified Linear units before being sent to the recurrent units. The temporal structure of the task is the same as shown in Figure 1A, except for the pre- and post-feedback periods, which were not implemented in the model since they are not of interest in the current study; the trial type was drawn randomly at the beginning of each trial during training. Each network was trained on a different random sequence of trials.

**Fig. 6:**
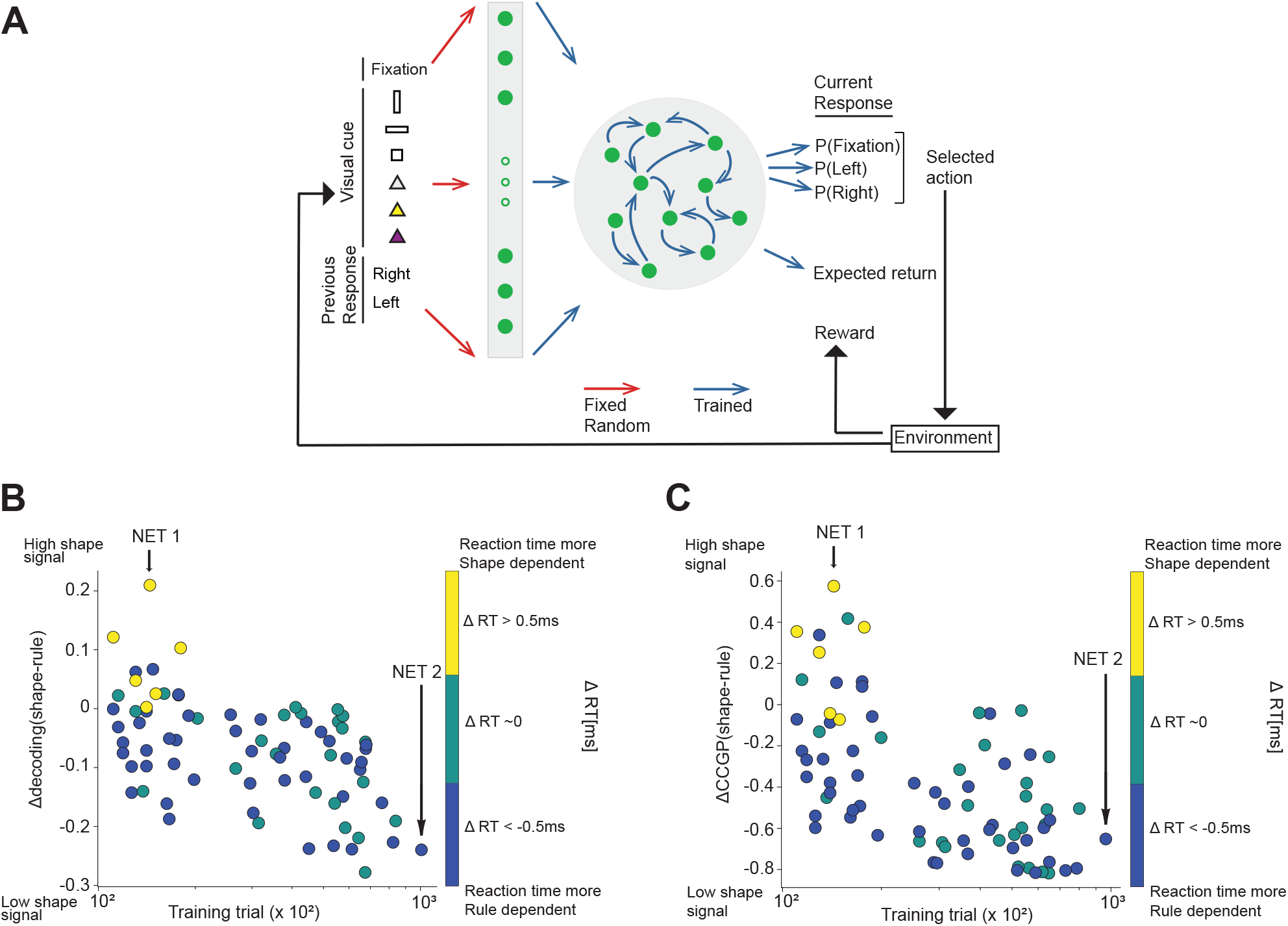
Studying individual diversity using 80 RNNs trained to perform the visually cued rule-based task. **A**) Architecture of the RNN, along with a schematic representation of the reinforcement learning setting (agent-environment interaction loop [36]). The inputs consist of one unit for fixation, six units for encoding visual cues (vertical rectangle, horizontal rectangle, or square combined with color, grey, yellow, or purple, denoted by triangles), and two units for encoding the previous response (right or left). The input is passed through an expansion layer of 100 Rectified Linear units. The output of the expansion layer is then processed by 100 recurrent units. At each time step, the network gives an output corresponding to fixate or select either right or left, represented by three units (network policy/actor), and the expected discounted return at that time step up to the end of the trial (value function/critic). **B**) Decoding results for all the recurrent networks, illustrating the difference in decoding accuracy of the shape and rule variables as a function of the required number of training trials to reach the performance criterion. The color bar indicates the difference in reaction time between shape and rule trials: positive values indicate a reaction time more influenced by shape irrespective of the rule, while negative values suggest a greater dependence on the rule regardless of the shape. A significant negative correlation is observed between the difference in decoding the two task variables and the number of training trials/amount of training (Pearson coefficient, *ρ* = −0.52, *p*-value = 10^−6^). This suggests that a higher amount of training required to reach the 90% performance threshold corresponds to greater decoding accuracy of the rule. Conversely, networks with a lower training requirement exhibit a stronger shape signal. The difference in decoding also shows a significant positive correlation with the difference in reaction time (Pearson coefficient, *ρ* = 0.35, *p*-value = 0.001), indicating that higher decoding accuracy for shape or rule corresponds to increased dependence of reaction time on the respective variable. The vertical black arrows indicate two example RNNs with high decoding accuracy for the shape (NET 1) and for the rule (NET 2). **C**) Similar to panel B, where the y-axis shows the difference between the CCGP for shape and rule. A significant negative correlation was observed between the difference in CCGP of the two task variables and the amount of training (Pearson coefficient, *ρ* = −0.48, *p*-value = 10^−5^), while a significant positive correlation was observed with the difference in reaction time (Pearson coefficient, *ρ* = 0.39, *p*-value = 10^−4^). 13

The output of the expansion layer was then processed by 100 recurrent units in a time-discretized vanilla RNN architecture, where the network provided outputs at each time step. The readouts included a scalar representing the temporal discounted expected return, defined as the output of the value function (critic), and a real vector with a length equal to the possible actions (actor policy). The action at each time step was determined by sampling from the categorical distribution obtained from the softmax of this vector (for examples of two trials, see Supplementary Figures 3A-B). We stopped the training when the network achieved at least 99% of complete trials and at least 90% of correct trials out of the complete ones on a validation batch of 10000 trials (Supplementary Figures 4A-B). See Methods for a detailed discussion of the model architecture, structure of the task, and the learning algorithm used to solve the task.

After terminating the training, we analyzed the activations/firing rates of the recurrent units on a separate testing set comprised of 10000 trials for each network. Our model aimed to explore the potential correlation between the representational geometries of shape and rule and the corresponding reaction times. To quantify this relationship, we first had to characterize the differences in the geometry across different RNNs. We defined a variable, Δ-decoding, which is the difference in the decoding accuracy of rule and shape during the cue presentation period. We also defined the variable Δ-CCGP, which is defined in the same way as Δ-decoding but for CCGP. These two variables provide a simple description of relevant differences in the representational geometry. Indeed, positive values of Δ-decoding indicate that shape is more strongly encoded than rule, resembling the representation observed in Monkey 1 (Figure 2A-B). Conversely, negative values indicated a stronger signal of rule compared to shape, resembling the representation observed in Monkey 2 (Figure 2C-D).

We evaluated the Δ-decoding (and CCGP) for each trained RNN and examined its correlation with the number of training trials needed by each network to reach the stop-learning criterion. Our analysis revealed a significant negative correlation between Δ-decoding and the training duration, as depicted in Figure 6B (Pearson coefficient, *ρ* = −0.52, *p*-value = 10^−6^). Similarly, the Δ-CCGP exhibited the same trend, as shown in Figure 6C (Pearson coefficient, *ρ* = −0.48, *p*-value = 10^−5^).

These findings suggest that the networks that reached the performance threshold with fewer training trials developed a stronger shape representation, as indicated by higher decoding accuracy (see Supplementary Figure 5). Interestingly, these networks also exhibited a larger CCGP for shape. The CCGP was generally high, indicating that these representations were abstract as in the experimental observations. Previous studies demonstrated that artificial simple feed-forward neural networks can easily generate this type of abstract representations using backpropagation and reinforcement learning algorithms [6, 31].

The networks with higher CCGP for the shape quickly found a policy to solve the task up to criterion while maintaining residual shape coding that comes from network initialization (the expanded inputs retain the low dimensional structure of the disentangled representations of shape and color). Conversely, the RNNs requiring more extensive training exhibited higher decoding accuracy and CCGP for the rule compared to the shape (Supplementary Figure 5). This result is not surprising because if the networks do not converge right away, the training is more extensive, and the representations tend to inherit the low dimensional structure of the output (see [31] for feed-forward networks), combining it with the structure in the inputs. These representations encode less strongly the shape, which is irrelevant for performing the task, and more strongly the rule, which is a combination of input features and their semantics. Thus, our results show that different RNNs, trained to solve the same task with high accuracy above 90%, exhibit distinct representational geometries for specific task variables, influenced by the number of training trials required to attain the performance threshold.

We then investigated the relation between the representational geometries of shape and rule with the reaction time: would the RNNs, exhibiting higher decoding accuracy and CCGP for shape, display a reaction time dependent on shape only? Analogously: would RNNs, with higher decoding accuracy and CCGP for the rule, exhibit a reaction time most strongly dependent on the rule? We compared the average difference in reaction time between trials with different shapes of visual cues to the average difference in reaction time between trials with different rules (see the definition 16 in Methods for details). Briefly, this comparison yielded a variable termed ΔRT, which was assigned to each RNN. Positive values of ΔRT indicated that, on average, reaction times were influenced by the identity of the shape, regardless of the rule. Conversely, negative values suggested that reaction times were influenced by the rule, regardless of the shape.

We discovered a significant positive correlation between Δ-decoding and ΔRT across all the RNNs (Figure 6B, Pearson coefficient, *ρ* = 0.35, *p*-value = 0.001). Similarly, a significant positive correlation was observed between Δ-CCGP and ΔRT (Figure 6C, Pearson coefficient, *ρ* = 0.39, *p*-value = 10^−4^). These results indicate that RNNs with a stronger shape signal during cue presentation typically require a shorter training period and exhibit average reaction times that change based on the identity of the shape, regardless of the rule. To gain further insights, we selected, among all the networks, the two ones with the highest shape and rule signals, respectively, and we analyzed them in detail. Specifically, we computed the decoding accuracy and CCGP over time from stimulus onset for the four uncorrelated task variables, as well as the behavioral performance and average reaction time.

Figure 7A illustrates the results for one of the examples RNNs, referred to as NET 1, which shows the highest shape decoding accuracy and CCGP, and it reaches the performance criterion with a small number of training trials among all the RNNs depicted in Figures 6B-C (see Supplementary Figure 6 for the decoding accuracy and CCGP along time for all the RNNs). During the presentation of the stimulus, the shape variable exhibits a high decoding accuracy, followed in time by the current response, previous response, and rule. The shape and previous response are also represented in an abstract format, with high CCGP, while the rule and previous response are not. Notably, although the rule and previous response can be decoded, they are not represented in an abstract format. Moreover, this network displays high behavioral performance for each task condition (*>* 90%, Figure 7B, left). Interestingly, the average reaction time varies depending on the shape of the cue, regardless of the rule. Specifically, the network responds faster to square cues compared to rectangle cues, irrespective of the rule (Figure 7B, right). Other networks that operate in the same regime also exhibit reaction times that depend only on the shape of the stimulus (not shown), but the RT for the squares is longer than for the rectangles. This is not surprising, given that the different visual input features (shape and color) are represented in the same way. It is possible that in the monkey brain, the colored stimuli elicit a more prominent response, and hence the reaction time would always be shorter for squares than for rectangles. This is something we can easily incorporate in the model, but we did not because we wanted to keep the model as simple as possible, and we found it interesting that we get the RT difference anyway, even when we do not introduce asymmetries in the input representations.

**Fig. 7:**
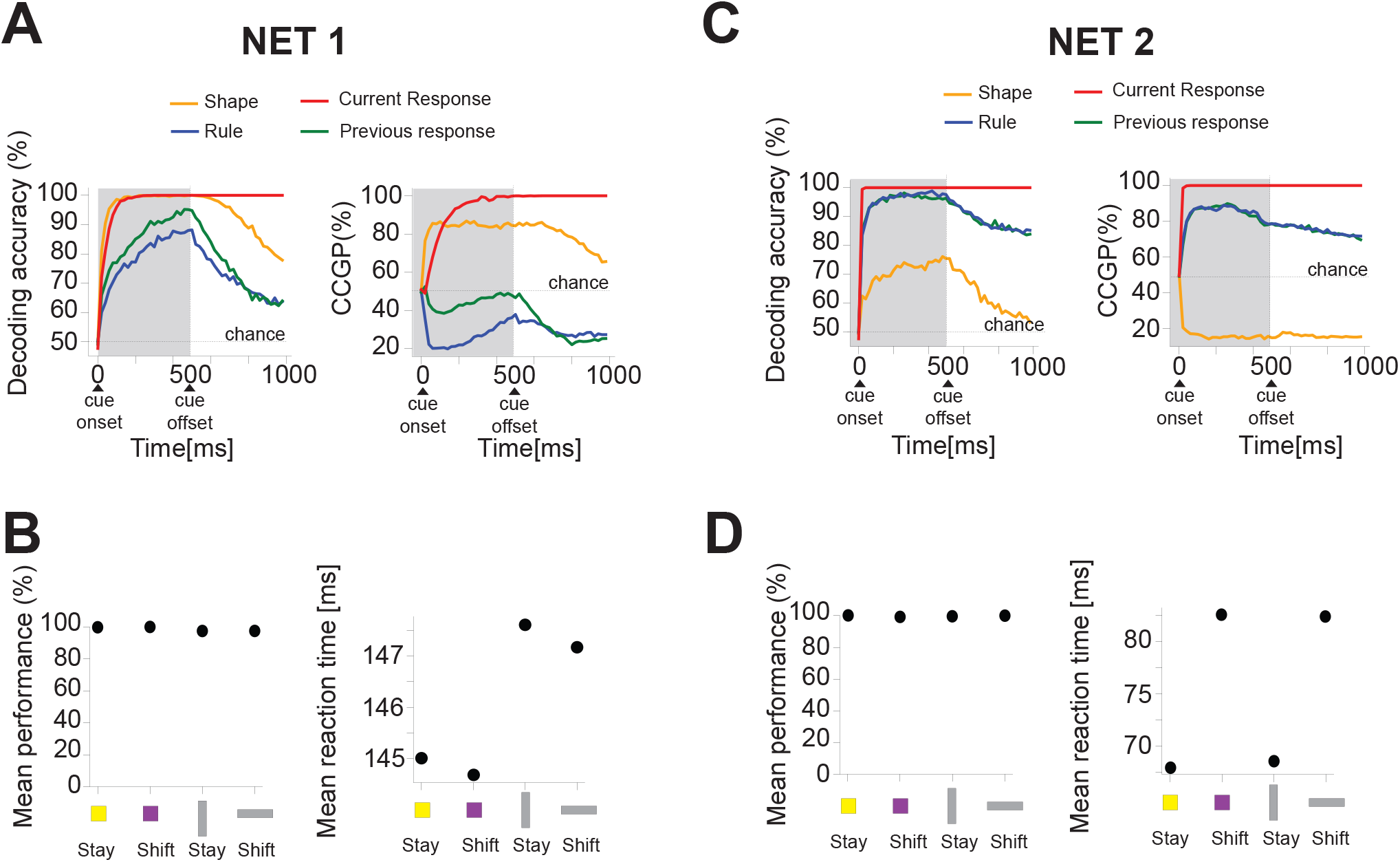
Representational geometries of task variables, behavioral performance, and reaction times of two example recurrent neural networks (RNNs). **A)** Left: decoding accuracy of the four main uncorrelated task variables for the RNN with high decoding accuracy for shape (NET 1 in Figure 6B). The activity is read out from the recurrent units. The shaded area indicates the period in which the stimulus is on the screen. After cue onset, the shape (orange line) is highly decoded, followed by the current response (red line), the previous response (green line), and the rule (blue line). Right: CCGP over time for the four main uncorrelated task variables. The shape (orange line), followed in time by the current response (red line), is represented in an abstract format after the cue presentation. **B)** Left: average behavioral performance for each of the four main task conditions displayed on the x-axis. Performance exceeds 90% for all conditions. Right: average reaction time for each of the four main task conditions. The reaction time of NET 1 exhibits dependence on the shape of the cue: on average, reaction time is faster for squares than for rectangles, irrespective of the rule. The representational geometry and reaction time patterns are similar to those of Monkey 1. **C)** Left: decoding accuracy of the four main uncorrelated task variables for the RNN with high decoding accuracy for the rule (NET 2 in Figure 6B). After cue onset, the rule (blue line) is highly decoded along with the current (red line) and previous (green line) responses. The shape (orange line) is decoded with lower accuracy compared to the other variables. Right: course over time of the CCGP for the four main uncorrelated task variables. The rule (blue line), current response (red line), and previous response (green line) are represented in an abstract format after cue presentation, while the shape is not (orange line). **D)** Left: average behavioral performance for each of the four main task conditions displayed on the x-axis. Performance exceeds 90% for all conditions. Right: average reaction time for each of the four main task conditions. The reaction time of NET 2 exhibits dependence on the rule: on average, reaction time is faster for the stay rule than for the shift rule, regardless of the shape of the cue. This RNN exhibits representational geometry and reaction time patterns similar to those of Monkey 2.

We performed the same set of analyses on the second example RNN, referred to as NET 2, which exhibits a high rule decoding accuracy and reaches the performance criterion with the highest amount of training trials among all the RNNs as shown in Figure 6B. During cue presentation, this network shows high decoding accuracy for the rule, previous and current response, which are also represented in an abstract format with high CCGP values. However, the shape variable is decoded with much lower accuracy and is not represented in an abstract format (Figure 7C). Similar to the previous example of NET 1, this network achieves high task performance across different task conditions (Figure 7D, left). Differently from the previous network example, the reaction time varies based on the rule, irrespective of the shape of the visual cue (Figure 7D, right). Specifically, it shows a shorter average reaction time for the stay rule compared to the shift rule, regardless of the shape.

One notable distinction between the model simulations and the actual data is in the representation of the previous response: in both models, it is abstract, whereas in both monkeys, it is not. This discrepancy can be attributed to our simplifying assumption that the previous response variable is entirely disentangled from the visual cue. Moreover, we provided the previous response as an additional input at the cue onset, while in the experiment, animals had to remember it from the previous trial. Another distinction between the models and actual data is the average RTs of the two example networks, which show different values (Figures 7B-D). Indeed, we did not fit the RTs in the models to reproduce the actual RTs of the animals. Our aim was to reproduce the patterns of RTs for different shapes and rules that emerged naturally from learning. Overall, we observed a huge variability in the average RTs of all the networks (Supplementary Figure 7), and it could eventually be rescaled in the model to match the data.

In summary, the representational geometries and reaction time patterns observed in the first example RNN (NET 1) resembled the neural and behavioral results observed in Monkey 1 (Figures 2A-B and Figures 5B-C). This network is probably operating in a lazy regime [32, 33, 34, 35], in which the neural representations inherited from the inputs are only slightly modified to perform the task correctly. Notice that shape is disentangled from the other task-relevant variables in the original inputs, and the random projections, despite being non-linear, only partially distort the geometry of the representations (see Supplementary Figure 8). This explanation is compatible with the observation that this network reaches high performance with more rapid training. Conversely, the results obtained from the second example network (NET 2) mirrored the neural and behavioral outcomes observed in Monkey 2 (Figures 2C-D and Figures 5D-E). High performance requires longer training (probably a rich regime), which enables the network to learn a representation that better reflects the task structure.

## Discussion

Traditionally, studies on the primate brain focused on the features of the recordings that are conserved across monkeys. It is uncommon to report and discuss differences between monkeys and other animals often because it is difficult to study and interpret them. Here, we showed that it is possible to find clear differences between the representational geometry of two monkeys and that they correlate with subtle but significant behavioral differences. One of the advantages of our approach, based on the analysis of the neural representational geometry, is that it allowed us to study systematically many different interpretable aspects of the geometry of the representation that potentially cause different behaviors. To characterize the representational geometry, we considered the decoding accuracy and the cross-condition generalization performance (CCGP) for every possible dichotomy of the experimental conditions. The number of dichotomies grows rapidly with the number of conditions, almost exponentially for balanced dichotomies (using Stirling approximation ∼ 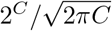 where *C* is the number of conditions). Even though some of the dichotomies are correlated (the full characterization of the geometry requires only ∼ *C*^2^ numbers), we can still systematically examine a large number of potentially different behaviors. Moreover, the dichotomies are interpretable and often correspond to some of the task key variables. This is the case in our analysis, in which the dichotomies with the highest decoding accuracy and CCGP during the cue presentation correspond to the shape of the visual stimulus for one monkey and the rule for the other. These dichotomies suggested a way to compute the reaction time for different groups of conditions and revealed significant differences in the behavior, which were not detectable from the initial analysis of the bare performance.

The analysis of the geometry revealed that there is an interesting “structure” in the arrangement of the points that represent different conditions in the firing rate space: for one monkey, the shape of the visual cue is an important variable (a more “visual” monkey), and for the other, it is the rule (a more “cognitive” monkey). This essentially means that for the first monkey, the points corresponding to different conditions in the firing rate space are grouped according to shape if one projects the activity on the coding direction of shape (notice that it is only in this subspace that the points cluster, as in the original space the points are still distinct and allow for the encoding of other variables). Analogously, the points are grouped according to the rule in the other more “cognitive” monkey. Both geometries and even the one in which the points are at random locations in the firing rate space (e.g. when the monkey is basically using a lookup table strategy, for which each visual cue is uniquely associated with a mapping from the previous response to the current response) allow for high performance. This is probably why we cannot see significant differences in the overall performance of the two monkeys. Moreover, the higher value of the shattering dimensionality in Monkey 1 than in Monkey 2 might support the hypothesis of the implementation of a more “visual” strategy to solve the task. However, these geometries have different computational properties that can only be revealed in novel tasks involving generalization or learning of new rules. For example, the more “cognitive” monkey, for which the rule is in an abstract format, would probably learn rapidly a novel task in which the rules are the same but the visual cues change. The new visual cues could be “linked” to the pre-existent groups that represent in an abstract format the two possible rules. The other more “visual” monkey is probably in a different learning stage, and the grouping, which is useless for performing the task or for generalizing to new similar tasks, is mostly dictated by the representations of the sensory inputs, as in the early stages of learning of the simulated network.

The simulation of Recurrent Neural Networks (RNNs) trained to perform the task used in the experiment with high accuracy revealed a significant correlation between the representational geometry of task variables and reaction times, and that the two monkeys could have gone through a different amount of training. Indeed, the models showed that those networks that reached the performance criterion with the smallest amount of training also have a higher signal for the shape over the rule, which resembles the representational geometry of Monkey 1. On the other hand, those networks that reached the threshold with more training developed a stronger signal for the rule than the shape, which resembles the representation in Monkey 2. Unfortunately, we do not have data collected during monkey training, but we know that, due to individual differences in the level of perseveration for which Monkey 2 tended to stay with the same response between consecutive trials, the training process was slightly different. In particular, Monkey 2 required a longer training than Monkey 1 to learn to switch between the stay and shift rule and establish the contingency between visual cues and the rule. This empirical note on the behavioral training of the two monkeys is in line with the correlation we observed between the representational geometry of the task variables and the amount of training across all the trained RNNs.

Moreover, with our models, we could reproduce the reaction time patterns, and we studied the relation between the representational geometry of the task variables and reaction time. The models with the highest signal for the shape of the visual cue showed a reaction time that, on average, significantly changes with the shape regardless of the rule, and it resembles the results we observed in Monkey 1. On the contrary, the models with the highest signal for the rule have a reaction time that, on average, significantly changes with the rule regardless of the shape, and it resembles the results we observed in Monkey 2.

It is possible that those networks that reached the performance threshold with a small number of training trials found an optimal policy to solve the task, probably due to the network initialization. Conversely, the RNNs requiring more extensive training developed a reduced representation of the shape, which in principle is useless to solve the task, in favor of a more robust rule representation. The use of different policies as a function of training amount, which is worth investigating in more detail for further studies, might reflect the development of different policies or strategies, also in the two monkeys to solve the same task, mirrored by different representational geometries of task variables correlating with different reaction times. For example, Tsuda et al. [37] recently showed that the different strategies of monkeys and humans in solving a working memory task (monkeys seem to apply a recency-based strategy while humans a target selective strategy [38, 39]) could correspond to two different learning stages of a simple recurrent neural network.

Although the model suggests that the differences are due to the training duration, it is also possible that the monkeys would have adopted different policies even at the same learning stage. We know from machine learning studies on curriculum learning that artificial neural networks can solve the same task in different ways depending on the order of presentation of the samples and, more generally, on the details of the learning process [40, 41]. Preliminary results on the RNNs indicate that an unbalanced distribution of trial types during the training phase can introduce biases in the resulting policies (data not included). Differences in strategies have been described in experimental studies, in particular in the information representations of the reward [42], in the strategies adopted by two monkeys to solve the same task [43], and in some abstraction tests [44]. A recent study, using a more complex task as the well know pac-man game, has even shown that different strategies can be flexibly switched based on different task demands [45].

In our study, we did not test whether abstract representations could lead to generalization to new stimuli. Introducing a generalization test would have allowed, for example, to test whether the abstract format of the rule in the second monkey generated a faster generalization to a new set of rule cues than in the first monkey. Future studies on abstraction should be planned to test whether the task variables encoded in an abstract form, as opposed to those that are not, would facilitate the generalization of the rules to new items or conditions.

The ability of generalization has been reported by several studies on macaques [46, 47, 48, 44]. For example, Falcone et al. [48] have shown that monkeys can transfer the nonmatch-to-goal rule from the object domain to the spatial domain in a single session, and Sampson et el. [44] have shown that abstraction can allow generalizing to new conditions, such as new foods, of the rule to choose the worst between two options.

Moving to chronic recordings surely offers the opportunity to follow in time the formation of neural representational geometries by recording before, and during the training phases, and after a task is fully learned. Planned behavioral generalization tests to new task conditions are critical to test the correlation between the geometry of the representation of a given variable and the animal performance in generalization tasks. These future studies will probably highlight even more individual differences and will allow us to define more precisely what a strategy is and how it is represented in the brain and to predict and test behavioral consequences in a number of novel situations.

## Author Contributions

A.G. and S.T. conceived and designed the experiments and collected the data. V.F, F.S., A.G., and S.F. conceptualized and developed the analyses. V.F. and F.S. analyzed the data under the supervision of S.F. V.F. and A.B. conceived and developed the model under the supervision of S.F. The data was interpreted by V.F., A.B., F.S., S.T., A.G., and S.F., who also wrote the article.

## Acknowledgments

We are grateful to L.F. Abbott for his comments on the manuscript and for insightful discussions. A.G. and S.T. thank S. Wise and A. Mitz for their numerous contributions. We thank M. Alleman, A. Fanthomme, R. Gulli, J. Johnston, J. Minxha, R. Nogueira, L. Posani, K. Rajan, and M. Rigotti for many valuable and knowledgeable discussions.

## Materials and Methods

### Subjects

All the details about the experiment are reported in the original article [24]. Here we give only a brief description of these details.

Two male rhesus monkeys (Macaca mulatta, 10–11*kg* in weight) were trained to perform a visually cued rule-based task. All experimental procedures were in agreement with the Guide for the Care and Use of Laboratory Animals and were approved by the National Institute of Mental Health Animal Care and Use Committee.

Each monkey, while performing the task, sat in a primate chair, with the head fixed in front of a video monitor 32*cm* away. An infrared oculometer (Arrington Research, Inc., Scottsdale, AZ) recorded the eye positions.

## Data collection and histology

Up to 16 platinum iridium electrodes (0.5–1.5*M* Ω at 1*kHz*) were inserted into the cortex with a multielectrode drive (Thomas Recording) to record single-cell activity from dorsolateral prefrontal cortex (Figure 1C). The recording chambers (18*mm* inner diameter) were positioned and angled according to magnetic resonance images (MRI). The single-cell potentials were isolated off-line (Off Line Sorter, Plexon), based on multiple criteria, including principal component analysis, the minimal interspike intervals, and close visual inspection of the entire waveforms for each cell. Eye position was recorded with an infrared oculometer (Arrington Research). The recording sites were localized by histological analysis and MRI (see Tsujimoto et al. [24] for more information).

## The behavioral task

A sequence of the task events of the visually cued rule-based task is shown in Figure 1A [24, 49, 50, 51]. For clarity, previous works’ authors referred to this task as the visually cued strategy task. The stay and the shift rules were designed as strategies because they represented a simplification of the repeat-stay and change-shift strategies used in previous neurophysiological studies [52, 53]. These two strategies were identified by Bussey et al. [54] studying the behavior of monkeys during the learning of visuomotor associations. The monkeys in their study spontaneously adopted the strategies to facilitate learning. As opposed to the previous studies of this task, here we refer to “strategy” as a possible way adopted by the monkey to solve the task, and to “rule” what is instructed to the monkey to perform the task. In each trial, the monkey was required to make a saccade towards one of the two spatial targets, according to a shift or stay rule cued by a visual instruction (Figure 1B). The appearance of a fixation point (a 0.6^*°*^ white circle) located at the center of the video screen, with 2 peripheral targets (2.0^*°*^ white square frames) placed 11.6^*°*^ to the left and right of the fixation point, represented the beginning of a trial. The monkey had to maintain fixation on the central spot for 1.5*s*; after that, a cue period of 0.5*s* followed. During the cue period, a visual cue appeared at the fixation point. In each trial, one visual cue was chosen pseudorandomly from a set of four visual cues: a vertical (light gray) or horizontal (light gray) rectangle with the same dimensions (1.0^*°*^ × 4.9^*°*^) and brightness, or a yellow or purple square with the same size (2.0^*°*^ × 2.0^*°*^) (Figure 1B). Each visual cue instructed either the stay or shift rule. The stay rule, instructed by the vertical rectangle or the yellow square, cued the monkey to choose the same target chosen in the previous trial (as shown in the two consecutive trials’ example in Figure 1A). Conversely, the horizontal rectangle or the purple square instructed the shift rule, which required the monkey to choose the target not chosen in the previous trial. The end of one trial and the beginning of the next one were separated by an intertrial interval of 1*s*. The first trial required a random choice of the target since no previous response could be integrated with the information on the current rule. Moreover, in the first trial, the monkey was always rewarded. The monkey had to maintain the fixation on the central point during the whole fixation period (1.5*s*) and the cue period (0.5*s*) as well as during a subsequent delay period of 1.0, 1.25, or 1.5*s*, pseudorandomly selected. The fixation window was a ±3^*°*^ square area centered on the fixation point. Both monkeys maintained fixation accurately and rarely made a saccade within the fixation window [24, 49]. Any fixation break during the fixation, cue, or delay periods led to abortion of the trial. The fixation point and the two peripheral targets were kept on the screen for the whole duration of the delay period. The disappearance of the fixation spot represented a go signal, instructing the monkey to choose one target by making a saccade to one of them. When the monkey fixated on one of the targets, both squares became filled. The entry of the gaze into the response window was labeled as target acquisition. The monkey had to maintain the fixation on the target for 0.5*s* (pre-feedback period). Any fixation break during the pre-feedback period led to abortion of the trial. After the pre-feedback period, in the case of correct response, feedback was provided as a liquid reward (0.2*ml* drop of fluid) or, in case of an incorrect response, as red squares over both targets. In the case of an error, the same cue was presented again in the following trial, called the correction trial. Correction trials were presented until the monkey responded correctly. Usually, after an error, there was not more than a correction trial [24, 49].

### Neurons and trials sample selection, pseudo-simultaneous population trials, and task conditions definition

We analyzed the neural activity of each monkey separately, only in complete and correct trials, from 400*ms* before the cue onset until 500*ms* after the cue offset. Linear decoders were trained and tested on pseudo-simultaneous population trials (pseudo trials). We defined a pseudo trial as the combination of spike counts randomly sampled from every neuron in a specific time bin and task condition [2]. The task condition is one of the eight possible combinations of task variables listed in Figure 1D. We analyzed the activity of neurons recorded in at least five trials per task condition.

Pseudo trials were generated as follows: given a time bin *t* and task condition *p*, for every neuron, we randomly picked a trial of task condition *p*, and we computed the spike count in the time bin *t*. The single pseudo trial *γ*, for condition *p* at time bin *t*, is then 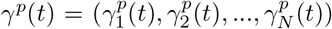, where *N* is the number of recorded neurons, and 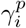 (*i* is the neuron identity, *i* = 1, …, *N*) is the spike count. We repeated this procedure 100 times, ending up with 100 pseudo trials per task condition and time bin.

Since we did not know a priori which task variables are represented by the neural ensemble, and in order not to introduce any bias in the selection of the task variables to decode, we defined a dichotomy as each pairing of the task conditions in a group of four, for a total of 35 dichotomies [6]. Each dichotomy is a variable that could be decoded. Four of the 35 dichotomies overlap with the task variables. All the other dichotomies cannot be explicitly interpreted in terms of any of the task variables, but rather as a combination of task variables which we referred to as other dichotomies. In particular, the four dichotomies that overlap with the task variables are the previous response, rule, current response, and the shape of the visual cue (Figure 1D). The latter identifies whether the visual cue was a rectangle or a square, which could also be interpreted as a grey-colored and non-grey-colored cue.

### Decoding of the neural population activity

For each dichotomy, which is a binary variable, we trained a Support Vector Machine (SVM) classifier with a linear kernel [55] to classify the spike count into either of the two values of the dichotomy. In all the SVM classifiers, we set a regularization term equal to 10^3^. We tried a range of regularization terms from 1 up to 10^3^, without any significant change in the final results. We decoded the neural activity in a 200*ms* time bin stepped by 20*ms* along time from 400*ms* before the cue onset until 500*ms* after the cue offset. The linear classifier was trained on pseudo trials built from randomly selected trials. In more detail, for every neuron, we selected 80% of the trials as a training set, and the remaining 20% as a testing set to build the pseudo trials. We cross-validated the linear decoder 100 times, by randomly choosing the 80% of the pseudo trials as training set, and the remaining 20% as testing set. We showed the final accuracy of the linear decoder as the ratio between the number of correct predictions to the total number of predictions on the testing set averaged across the cross-validations. To evaluate the statistical significance of the neural signal, we built a null model by randomly shuffling the task condition labels among the pseudo trials. For each shuffle, we trained a linear decoder on the shuffled training set, and we assessed its accuracy on the shuffled testing set. We repeated the shuffle procedure 100 times, obtaining a null model distribution. We defined the chance interval as the interval between 2 standard deviations of the null model distribution around the chance level at 50%.

The data were extracted by custom MatLab functions (The MathWorks, Inc., Natick, MA, USA). All decoding analyses were performed by using scripts of the scikitlearn SVC package along with custom Python scripts [55].

### Neural representation of variables in an abstract format and the Cross Condition Generalization Performance

After assessing which task variables are decoded, we asked in what format they are represented. In particular, we asked whether they are represented in an abstract format. A variable could be defined to be in an abstract format when a linear decoder trained to classify the value of the variable can generalize to new task conditions never used for training. To assess to what extent a variable is in an abstract format, we computed the Cross Condition Generalization Performance (CCGP), that is, the performance of a linear decoder in generalizing to new task conditions not previously used for training [6]. The difference between the traditional cross-validated linear decoder and the cross-condition generalization is in the data used for training and testing the classifier. In the traditional-fashioned decoding analyses, a decoder is trained on a sub-sample of trials randomly picked from each (experimental) condition, and tested on the held-out trials retained from each condition. In the end, the decoder is trained and tested on all the conditions, and the generalization is only across trials. The CCGP, instead, is computed by training a linear decoder only on a fraction of trials from a subset of conditions and tested on trials belonging to new conditions not used for training. The generalization is now not only across trials but also across conditions.

We assessed the CCGP for each of the 35 dichotomies as follows. Given a dichotomy, defined as a pairing of task conditions in a group of four, we trained the decoder to classify the value of the dichotomy using trials from three task conditions from each side of the dichotomy and tested it on the one held-out condition from each side. Since each side of the dichotomy has four task conditions, there are 16 possible ways of choosing the training and testing condition set. For each choice of training and testing set, we applied 10 cross-validations, randomly choosing 80% of training trials and 20% of testing trials. We reported the average performance across all 16 possible choices of training and testing conditions and the 10 cross-validations for each dichotomy. To assess the statistical significance of the CCGP, we built a null model where the geometrical structure in the data was destroyed while keeping the variables still decodable [6]. To do that, we applied a discrete rotation to the noise clouds (the trials firing rate of each condition) by permuting the axes of the firing rate space and randomly assigning neural activity to neurons. We repeated this procedure for each cluster separately. We generated 100 null models, and for each of them, we computed the CCGP for all dichotomies, as done on real data. We defined the chance interval for the CCGP measure as the interval between 2 standard deviations of the null model distribution around the chance level at 50%.

### Multi Dimensional Scaling analysis

We used the Multi-Dimensional Scaling (MDS) transformation to seek a low-dimensional representation of the data. We computed the metric MDS, where the dissimilarity matrix was built as follows. We averaged the neural activity, in a fixed time bin, across pseudo trials within each task condition, and we constructed a *p*_*c*_ *×p*_*c*_ matrix (with *p*_*c*_ indicating the number of conditions) which stored the Euclidean distance between the average firing rate between each paired condition. In order to keep information regarding the noise cloud of each task condition, we normalized the Euclidean distance matrix by the squared root of the sum of the variance of each condition along the distance direction between the two clouds. For the analysis based on a single pseudo trial (Figure 3B), the dissimilarity matrix was defined as a *p*_*t*_ × *p*_*t*_ matrix, with *p*_*t*_ indicating the total number of pseudo trials across all conditions. This dissimilarity matrix stored the Euclidean distance between the firing rate of each pair of pseudo trials, and it was normalized as described above.

### Behavioral analyses

We computed the behavioral performance and reaction times of each monkey separately, combining all the sessions we considered for the neural analyses. We computed the reaction time (RT) only in complete and correct trials. The RT is defined as the time difference between the go signal and target acquisition in each trial. In order not to bias the results due to outliers, we removed those trials with RT larger than 3 standard deviations from the mean. Since the neural analyses revealed that the difference between the two monkeys comes from different representational geometry of the rule and the shape of the visual cue, we grouped trials per rule (stay-shift) and shape (rectangle-square), for a total of four conditions. We compared the distribution of RTs of trials with different rules and shapes, separately. To test whether the RTs distributions were significantly different, we ran the Mann-Whitney U-test (p-value*<*0.05).

Moreover, for each of the previous four task conditions, we computed the average performance across the sessions. The error bar of the estimated average performance was assessed by applying the following formula [56]:

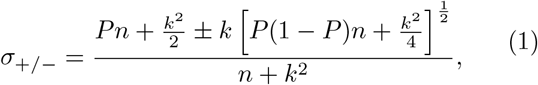

where *n* is the number of trials used to compute the performance across sessions, *P* is the average performance, and *k* is the confidence level in terms of standard deviation that we fixed equal to 2. To assess whether the performance was statistically different between different conditions, we applied the chi-squared test (p-value*<*0.05).

### Multi-Linear Regression Model for behavior

We fitted a multi-linear regression model on a single-trial basis to better investigate the behavioral differences between the two monkeys. We included only complete and correct trials in the model, and we discarded those trials with reaction times larger than 3 standard deviations from the mean as done in the behavioral analysis. For each trial, we took three independent binary input factors to the model: rule (+1/−1), previous response (+1/−1), and shape (+1/−1). We also included all the interaction terms. The output of the model is the reaction time, and the multi-linear model is defined as follows:

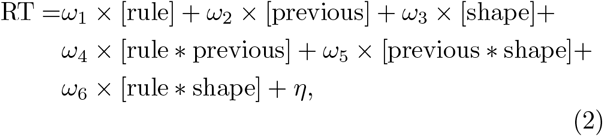

where *ω*_1,…,6_ are the weights of each factor, and *η* is a constant term. We fitted 100 models, each time randomly subsampling trials from each task condition, in each monkey separately. The number of trials per task condition was set to the minimum number of trials across conditions. We fitted each model by using the ordinary least squares method [57]. We compared the weights’ distributions across models between the two monkeys, for each factor, using the Mann-Whitney U test (p-value*<*0.05).

### Architecture of the Recurrent Neural Network Model

We trained 80 vanilla Recurrent Neural Networks (RNNs) to perform the visually cued rule-based task (see Figure 6A). Through random fixed weights *W* ^*rand*^, *N*_*u*_ = 9 input units are fully connected to an expansion layer of *M* = 100 rectified linear units (ReLU). The output from the expansion layer is passed through the input weights *W* ^*in*^ to *N* = 100 recurrent units. *W* ^*rec*^ defines the recurrent weight matrix (*N* × *N*). The readouts of the RNN are a single scalar representing the temporal discounted expected return (value function/critic), and a real vector with a length equal to the total number of possible actions (*N*_*a*_ = 3), which are fixation, right or left response (policy/actor). The final action was determined by sampling from the softmax distribution of this vector at each time step.

The input to the network, **u**(*t*)^*task*^, which is the input vector to the expansion layer, is defined as follows:

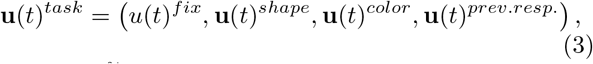

where *u*(*t*)^*fix*^ is a scalar that is equal to 1 when the network has to fixate, and it is set to 0 when the network is required to provide a response after the delay period; **u**(*t*)^*shape*^ is a one-hot vector of three units encoding the shape of the visual cue (horizontal, vertical rectangle, or square); **u**(*t*)^*color*^ is a one-hot vector of three units encoding the color of the visual cue (grey, yellow, or purple); **u**(*t*)^*prev*.*resp*.^ is a one-hot vector of two units encoding the previous response (right or left). This is a simplified version of the input with respect to the monkeys’ behavioral task where the animals had to retrieve the previous response from the earlier trial (here, we randomly sampled the previous response at the beginning of each trial because we reset the network at the end of each trial for simplicity, and we provided it to the network at the cue onset). For the current model, the input vector **u**(*t*)^*task*^ was randomly selected at the beginning of each trial so that the possible trial types were uniformly randomly sampled.

We defined a positive activity input **u**(*t*) to the recurrent units as [58]:

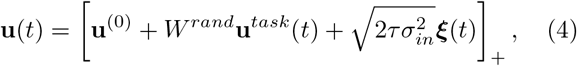

where *u*^(0)^ = 0.2 is a constant baseline term (equal for all the units), *W* ^*rand*^ are random weights from the nine input units *N*_*u*_ to the expansion layer units *M, τ* = 100*ms* is the neuronal constant, *σ*_*in*_ = 0.01 is the strength of the input noise, and ***ξ***(*t*) is Gaussian white noise with zero mean and unit variance, sampled i.i.d. across units and time. The ReLU non-linearity function [x]_+_ = max(0, x) maps the input currents to positive firing rates. The random weights *W* ^*rand*^ were sampled from a Gaussian with zero mean and standard deviation 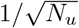, and they were kept fixed during the whole training phase. We can rewrite the Eq.4 in the discrete-time description with time step Δ*t* = 20*ms*, using the first-order Euler approximation, in the following way:

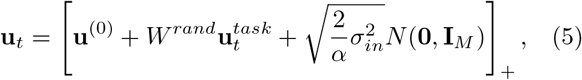

where *α* = Δ*t/τ*, and *N* (**0, I**_*M*_) is the multivariate Gaussian centered in zero with the identity as covariance matrix (of size *M* × *M*).

We described the *N*-dimensional recurrent units activity **r**(*t*) by the following dynamical equation [59]:

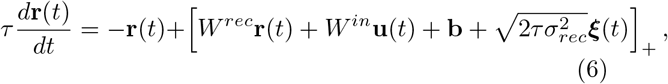

which, in a discrete-time formulation using first-order Euler approximation, becomes:

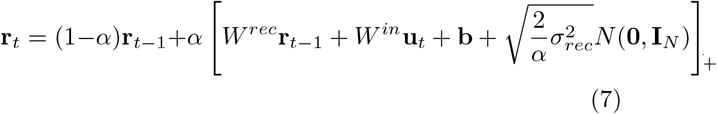

where *W* ^*rec*^ are the recurrent weights, *W* ^*in*^ are the initial weights from the expansion layer to the recurrent units, **b** is the bias term, and *σ*_*rec*_ = 0.05 is the strength of the recurrent noise. We initialized the recurrent connection weights *W* ^*rec*^ as a scaled identity matrix 0.5 *×* **I**_*N*_, where **I**_*N*_ is the identity matrix of dimension *N* × *N, N* = 100 recurrent units. The input weights *W* ^*in*^ connecting the *M* = 100 expansion layer units to the *N* = 100 recurrent units were initialized by sampling them from a Gaussian distribu<u>ti</u>on of mean zero and standard deviation equal to 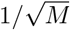. The bias **b** term was initialized to zero.

One of the output readouts of the RNN is the scalar representing the temporal discounted expected return *V*_*t*_ (value function/critic) at time *t*, defined as follows:

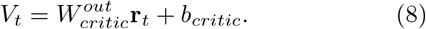

The second readout that implements the policy (actor) is a real vector of 3 units, each representing a possible action. The final action was determined by sampling from the softmax distribution of this vector as follows:

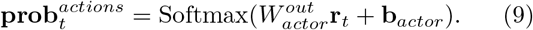

The output weight matrices 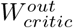 and 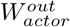 were both initialized by sampling from a Gaussian distribution of zero mean and standard deviation equal to 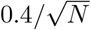. The bias terms *b*_*critic*_ and **b**_*actor*_ were initialized to zero.

For each network, we trained the following parameters: 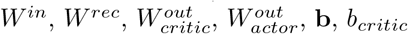 and **b**_*actor*_.

### Training of the RNN through the Proximal-Policy-Optimization

We trained the RNNs to perform the visually cued rule-based task using Proximal-Policy-Optimization (PPO), a state-of-the-art deep reinforcement learning algorithm [27]. We defined the Loss Function ℒ to be maximized on every training batch of trials, as a weighted sum of the PPO policy loss ℒ^*PPO*^, the state value function loss ℒ^*V F*^, and the entropy regularization term *S* as follows:

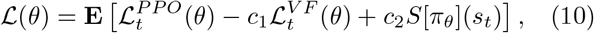

where *c*_1_ = 0.5 and *c*_2_ = 0.01 are hyperparameters determining the weights of the value function loss and the entropy regularization term, respectively. **E** represents the mean over a batch of training trials that is composed by unrolling 20 environments, simulated in parallel with the same agent, for *T* = 128 steps. This batch was then split into 4 mini-batches used for the optimization. *θ* refers to the collections of all the trained parameters, and *π*_*θ*_ is the policy used to sample the actions of the network given an input.

The policy loss ℒ^*PPO*^ is defined as follows:

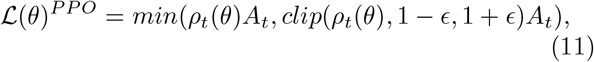

where 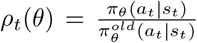 is the probability ratio of the current and old (before the update) policies, *ϵ* is a hyper-parameter set to 0.1, and *A*_*t*_ is the advantage function (similar to reward-prediction-error) defined as:

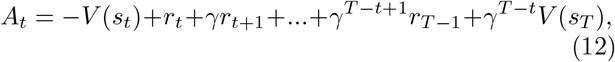

where the state *s*_*t*_ = **u**_*t*_ is the input to the network over time as defined in Eq. 5, *r*_*t*_ is the actual reward at time step *t, γ* is a standard temporal discount factor set to 0.99, and *V* (*s*_*t*_) is the value function computed at the state *s*_*t*_. The goal is to get a policy that maximizes future rewards in the interaction agent/environment loop.

The value function *V* (*s*_*t*_) is optimized in a supervised way by minimizing the following mean square error loss:

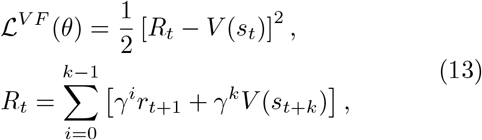

where *R*_*t*_ is the *n*-step bootstrapped discounted return at time *t, V* (*s*_*t*_) is the value function whose output is the expected return from state *s*_*t*_, *γ* is the discounted factor, *k* is the number of steps until the next state and it is upper bounded by the maximum unroll length *T*, and [*R*_*t*_ − *V* (*s*_*t*_)] is the temporal-difference error that provides an estimate of the advantage function for actor-critic.

Finally, the entropy regularization term that helps with exploration over exploitation is defined as follows:

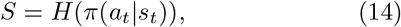

where *a*_*t*_, and *s*_*t*_, are the action, and the state, respectively, and *H*(*π*) is the entropy of the policy.

All the parameters were updated via gradient ascent and backpropagation through time using the Adam optimizer with default parameters [60]. In order to deal with the issue of the exploding gradients, we clip the gradient norm always to be ≤ 1.

### Visually cued rule-based task structure for the network model

We trained the networks to perform the visually cued ruled-based task, which is equivalent to the task that the monkeys were trained on, but without the feedback period (Figure 1A). The network is reset at the beginning of each trial with initial firing rate **r**_0_ = **0**. Each trial starts with a fixation period of *t*_*fix*_ = 1500*ms*. The time increment after each step is set to Δ*t* = 20*ms*. During the fixation period, the only non-zero input to the network is the fixation input, and the network has to choose the fixation action to take a reward of 0; otherwise, if it takes a right or left action, the trial is aborted, and a reward of −1 is issued.

After the fixation period, there is the appearance of one of the visual cues (see Figure 1B), which is randomly sampled at each trial, and it remains on for *t*_*cue*_ = 500*ms*. The fixation input is still active during the cue onset, and the network must continue to maintain fixation. The previous response is provided along with the visual cue. It is randomly sampled in each trial, and it is presented only during the cue period to reproduce the results we showed in Figure 2, where the previous response is mainly decoded around the visual cue presentation.

After the cue period, a delay period of *t*_*delay*_ of 1000, 1200, or 1500*ms* is randomly chosen in each trial, as in the monkeys’ task [61][24][49], where the only input to the network is the fixation. Again, the network can only choose to maintain fixation, and any other action results in punishment and abortion of the trial.

Subsequently, after the delay period, the fixation input is turned off, representing the go cue, where the network has a maximum of *t*_*dec*_ = 1500*ms* to make a decision, left or right. In this phase, if the network continues to hold fixation for more than *t*_*dec*_, the trial is aborted, and a reward of −1 is issued. If, on the other hand, the network provides the correct action (right or left), a reward of +1 is given, and the trial is terminated. However, if the network makes the wrong choice, it is punished with a reward of −1, and the trial is terminated.

The reaction time (RT) is calculated as the difference between the first time step the network makes a choice, right or left, *t*_*fin*_, and the time step of the go cue, *t*_*go*_: RT= *t*_*fin*_ − *t*_*go*_, where *t*_*go*_ = *t*_*fix*_ + *t*_*cue*_ + *t*_*delay*_. See Supplementary Figure 3 for 2 examples of correct trials after successfully training a model.

### Analysis of the correlation between the representational geometry of shape and rule with reaction times in RNNs

Since one main difference in the two monkeys, performing the same task with high accuracy, concerns the representational geometries of the shape of the visual cue and the rule during the cue presentation, we defined two variables, Δ-decoding and Δ-CCGP defined as follows:

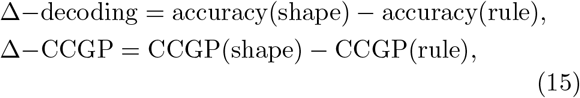

where accuracy(shape) and accuracy(rule) are the accuracies of the linear decoder in classifying the shape and rule, respectively, during the visual cue presentation, and analogously for the CCGP. We computed these variables for each of the RNNs. They are good indices that briefly summarize the difference in representational geometries of the shape and rule across all the RNNs. Indeed, positive values of Δ-decoding, or Δ-CCGP, suggest a stronger signal of the shape compared to the rule, resembling the representation observed in Monkey 1. Conversely, negative values indicatea stronger signal of the rule compared to the shape, resembling the representation observed in Monkey 2.

The second main result we obtained from neural data is the correlation of the representational geometries with reaction times (RTs). To assess the difference in reaction times, we defined a new variable ΔRT as follows:

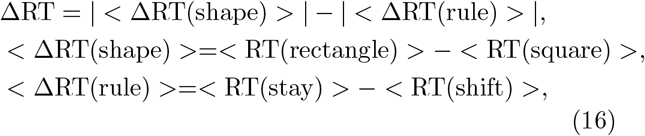

where *<*RT(rectangle)*>, <*RT(square)*>*, and *<*RT(stay)*>, <*RT(shift)*>*, are the average reaction time across trials with rectangle-shaped or square-shaped cues, and with stay or shift rules, respectively. We then computed the difference between the two averages and took the absolute value.

We assessed the ΔRT for each RNN: positive values of ΔRT indicate that, on average, reaction times are influenced more by the identity of the shape than the rule. Conversely, negative values suggest that reaction times depend more on the rule than on shape.

We subsequently correlated the Δ-decoding, or CCGP, with the number of training trials required to reach the performance threshold of at least 99% of complete trials and 90% of correct trials to stop the training, and with ΔRT, separately.

## Supplementary Figures

**Fig. 1:**
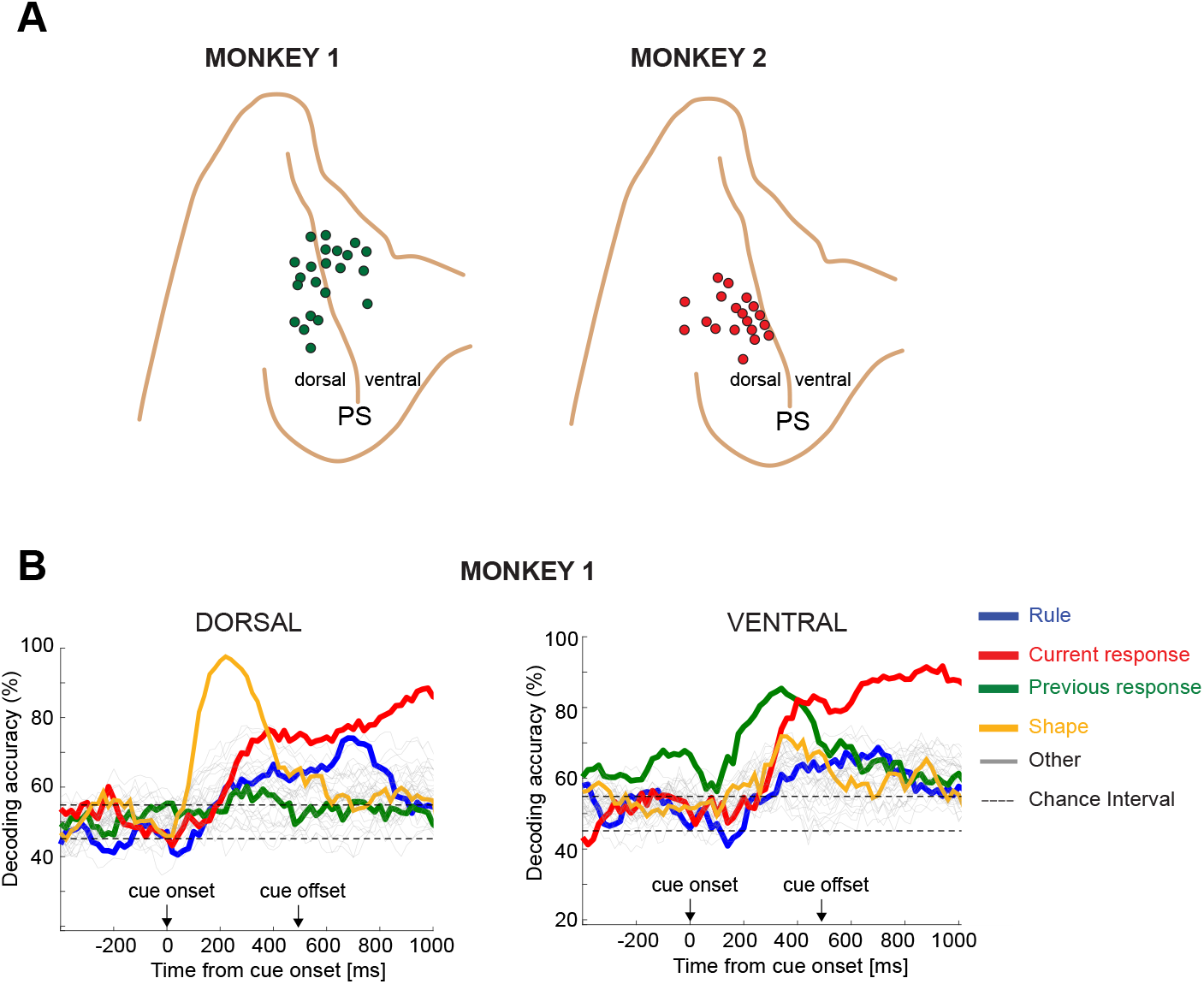
Recording sites for Monkey 1 and Monkey 2 in the dorsolateral prefrontal cortex (PFdl). **A)** Recording sites in PFdl for the two monkeys. **B)** Decoding accuracies of all dichotomies for Monkey 1 after splitting neurons in dorsal (106 neurons) and ventral (99 neurons) recordings with respect to the principal sulcus. PS: principal sulcus.

**Fig. 2:**
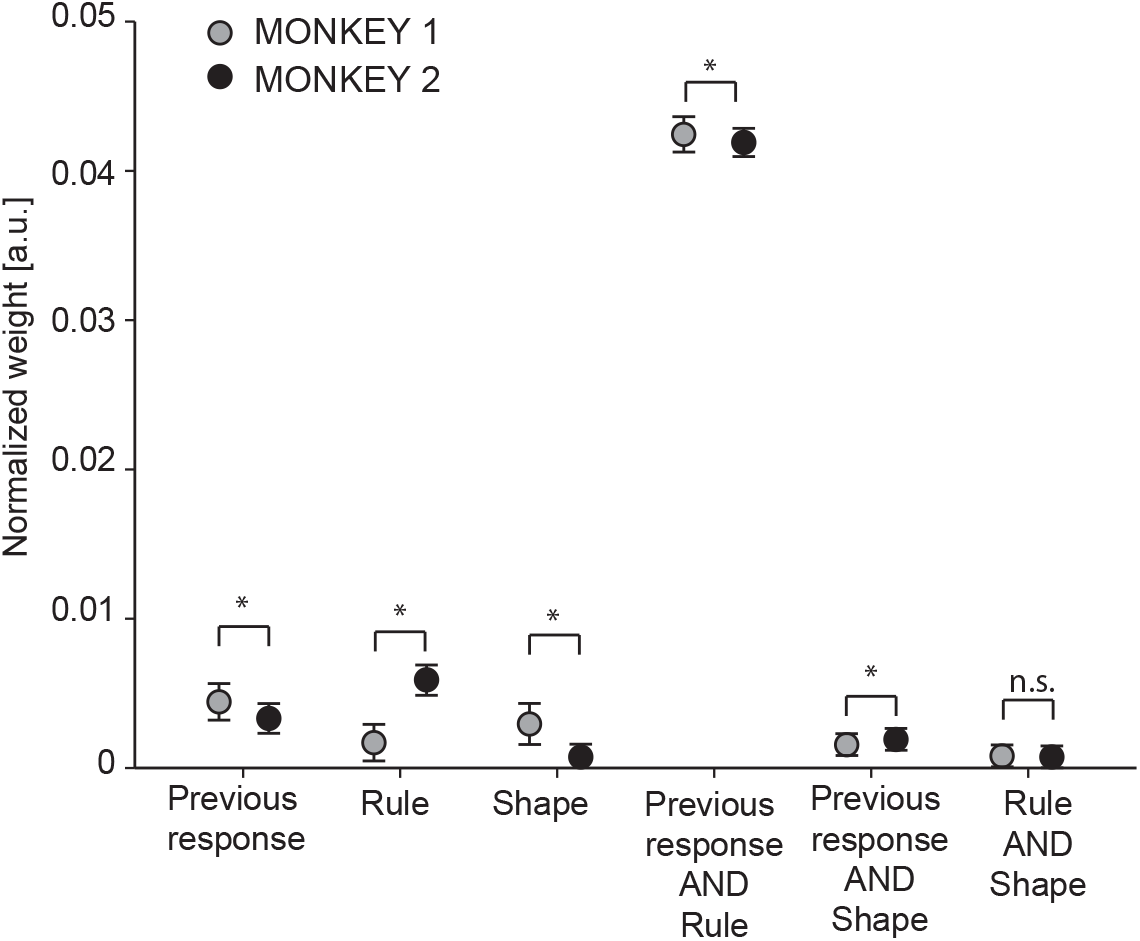
Multi-Linear regression analysis results. Mean of the distribution of the weights of 100 multi-linear regression models. The reaction time is predicted on a single trial using three factors: previous response, rule, and shape along with the interaction terms. The interaction term of the previous response with the rule is the strongest factor in both monkeys since the combination of these two variables is essential to elaborate the correct response. The error bars are the 2 standard deviations of weights across 100 models. n.s.: not significant; * Mann-Whitney U test: p*<*0.05.

**Fig. 3:**
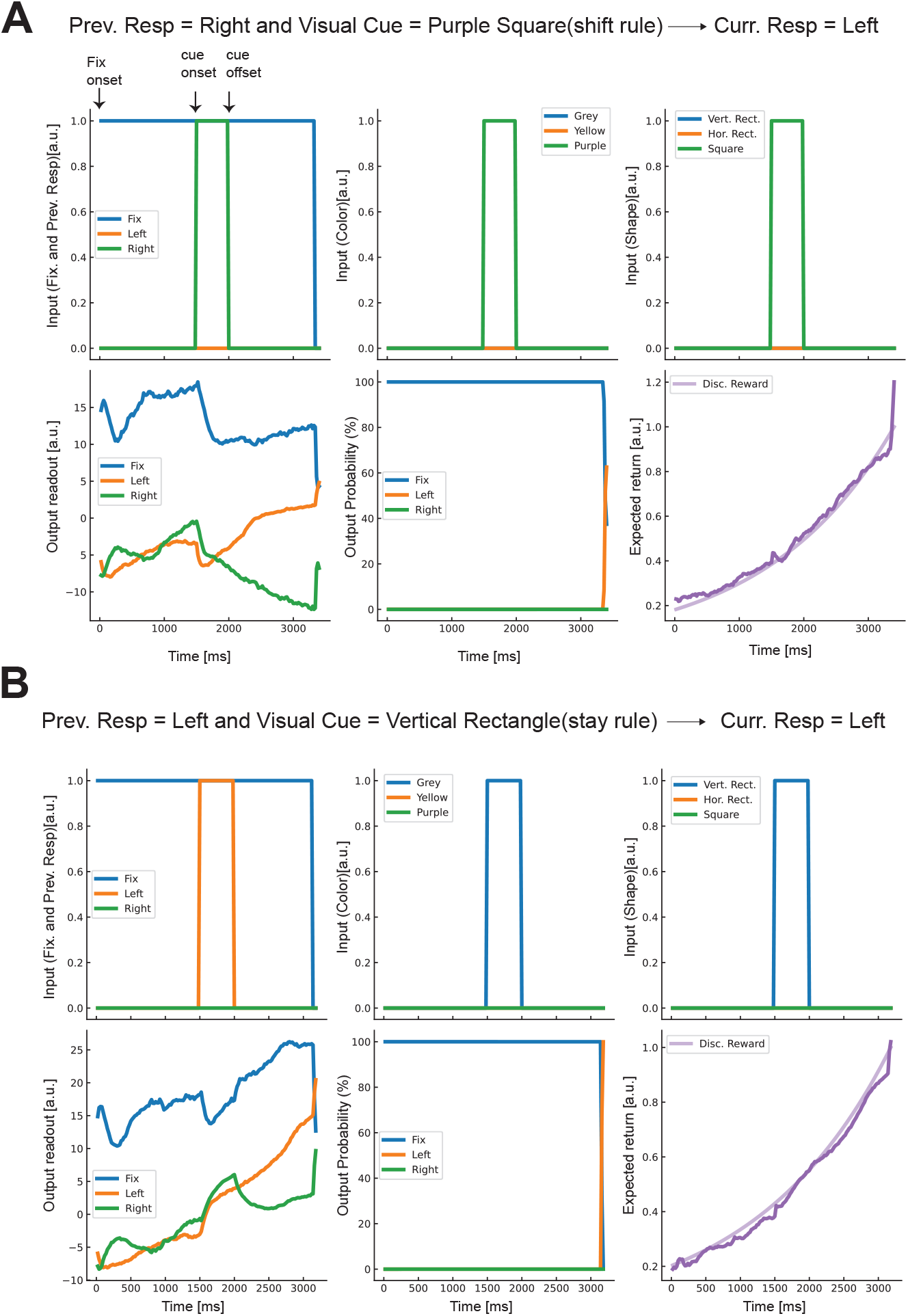
Example of two trials solved by an RNN. A). Example of a shift trial with the purple square visual cue and previous response right. The correct current response is left. The top row illustrates the three inputs to the network during the trial. Top-left: The fixation input (blue line) is constantly active before the go cue onset, while the previous response (green line) is given during the visual cue presentation for 500*ms*. This is a simplified version of the real task where the animal is required to retrieve the previous response from the previous trial. Top-middle: color input (purple) of the visual cue that, in this example, is the purple square. The purple signal is presented during the cue presentation, and it lasted for the 500*ms*. Top-right: the shape input (square) of the visual cue that in this example is the purple square. The bottom row illustrates the different output readouts from the network during the trial. Bottom-left: example of the policy output signal during time. The fixation output signal is high while the fixation input is on (blue line). After the presentation of the visual cue, the left response signal (orange line) starts to increase over the right signal (green line) and the fixation signal. Bottom-middle: output probability computed from a softmax of the policy output signals at each time step during the trial. Bottom-right: Expected return is defined as the value function output along the trial: it decays exponentially backward in time because of the temporal discounting imposed during training. **B)** Example of a stay trial. This is the same as in A, with the left previous response, along with the vertical grey rectangle visual cue, requiring left as the current correct response.

**Fig. 4:**
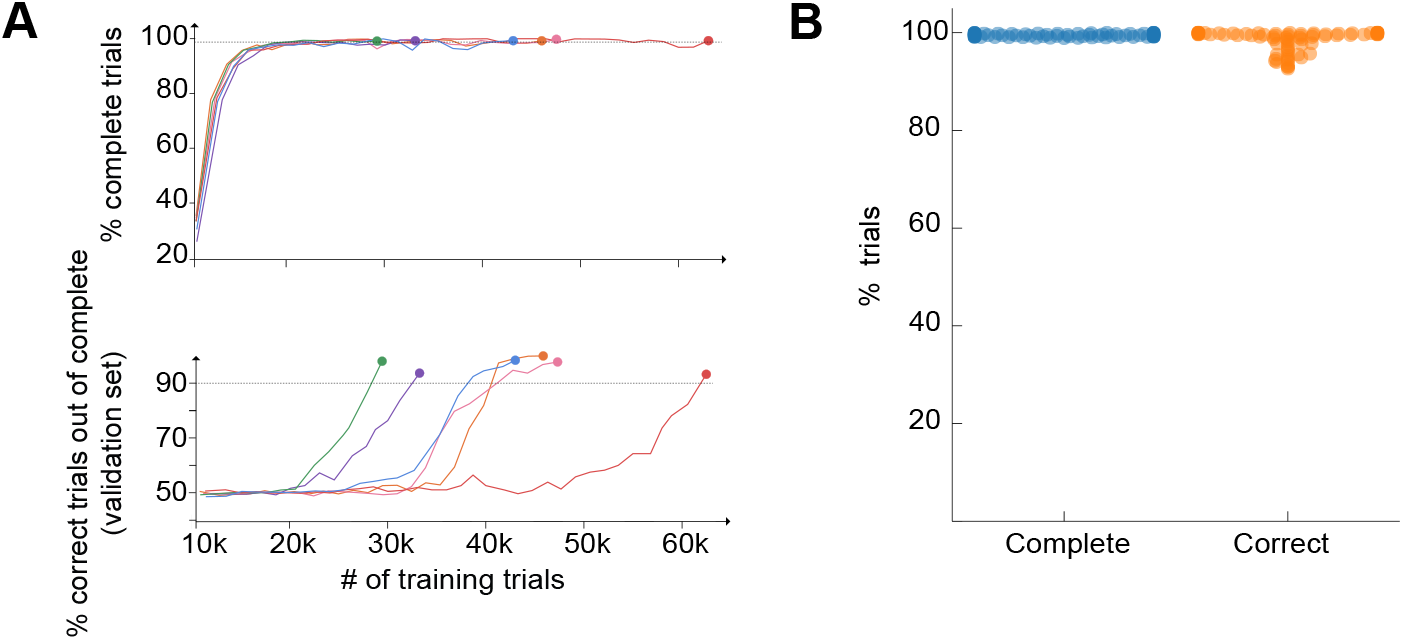
Percentage of complete and correct trials over complete trials of RNNs during training. **A)** Top: Percentage of complete trials of six example RNNs as a function of the number of training trials. The horizontal dashed line indicates the threshold of 99% of complete trials where we stopped the training. Due to the heterogeneity, some networks converged earlier than others, potentially due to differences in initial random conditions and random pattern presentation of trial types during training. The percentage is calculated after a fixed number of time steps over a batch of 10000 validation trials (see Methods). Bottom: Percentage of correct trials out of the complete trials shown above for the same six RNNs. The horizontal dashed line indicates the threshold of 90% of correct trials where we stopped the training. The performance is assessed on a validation set not previously used for training. The high level of idiosyncrasy is evident among RNNs converging to the threshold at different amounts of training trials. **B)** Percentage of complete and correct of the complete trials for each of the 80 RNNs after training. All the RNNs satisfy the convergence criterion, i.e., at least 99% of complete trials (left) and at least 90% of correct trials out of the complete trials (right). The percentage is calculated over a batch of 10000 testing trials for each network.

**Fig. 5:**
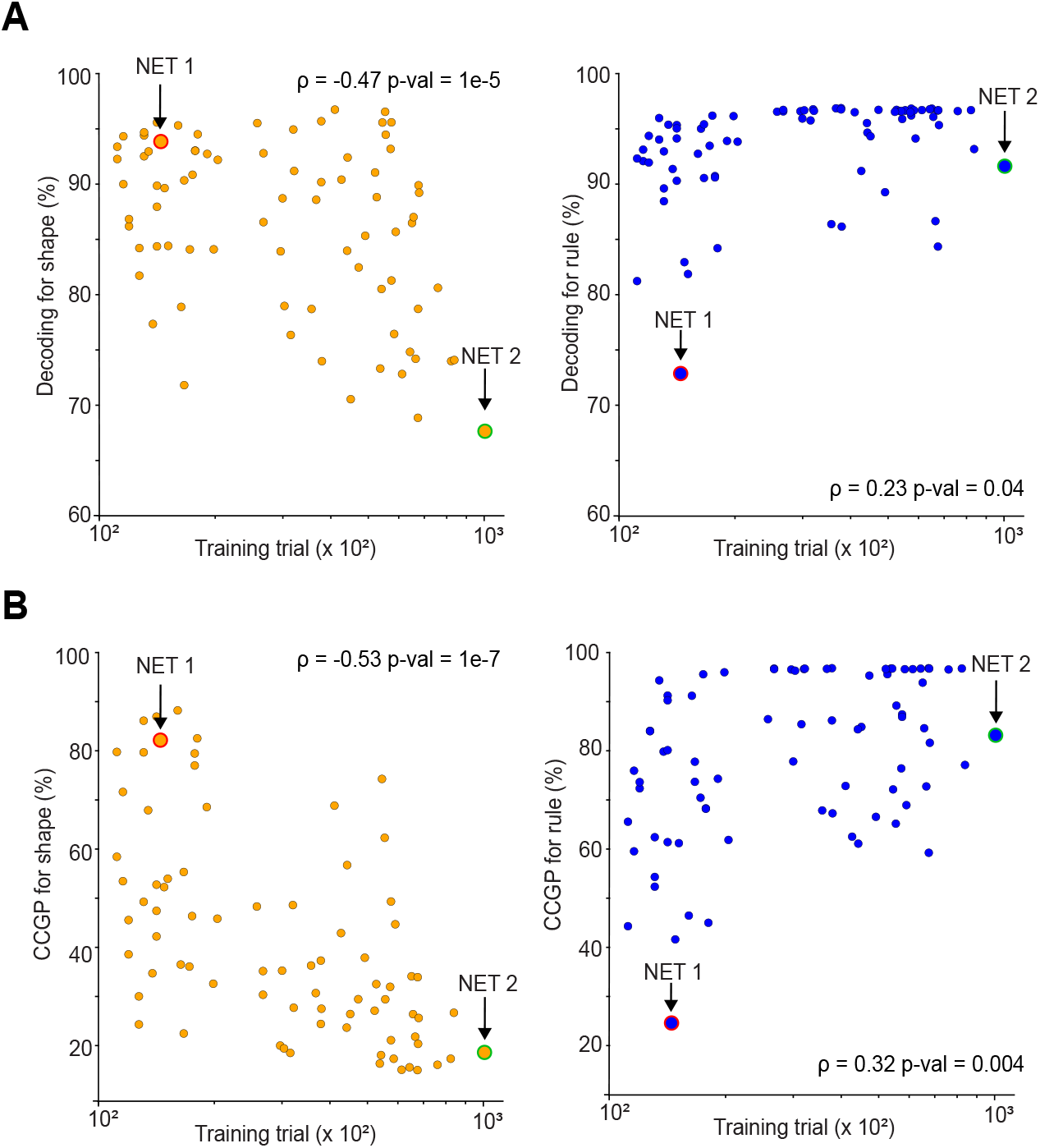
Decoding performance and CCGP for shape and rule during the cue presentation versus the number of training trials. **A)** Left: Decoding accuracy of the shape during the 500*ms* of cue presentation as a function of the number of training trials required to reach the 90% of performance threshold. The higher the amount of training, the lower the decoding accuracy for the shape (Pearson correlation, *ρ*=-0.47, p-value=1e^−^5). We enhanced the two example networks, NET 1 and NET 2, which resemble the monkeys’ shape and rule representations (see Figure 7). Right: Decoding accuracy of the rule as a function of the training amount. The higher the training amount, the higher the decoding accuracy of the rule (Pearson correlation, *ρ*=0.23, p-value=0.04). **B)** Left: CCGP for the shape as a function of the amount of training. As for the decoding accuracy, the higher the amount of training, the lower the CCGP (Pearson correlation, *ρ*=-0.53, p-value=1e^−^7). Right: CCGP for the rule as a function of the amount of training. The higher the amount of training, the higher the CCGP for the rule (Pearson correlation, *ρ*=0.32, p-value=0.004).

**Fig. 6:**
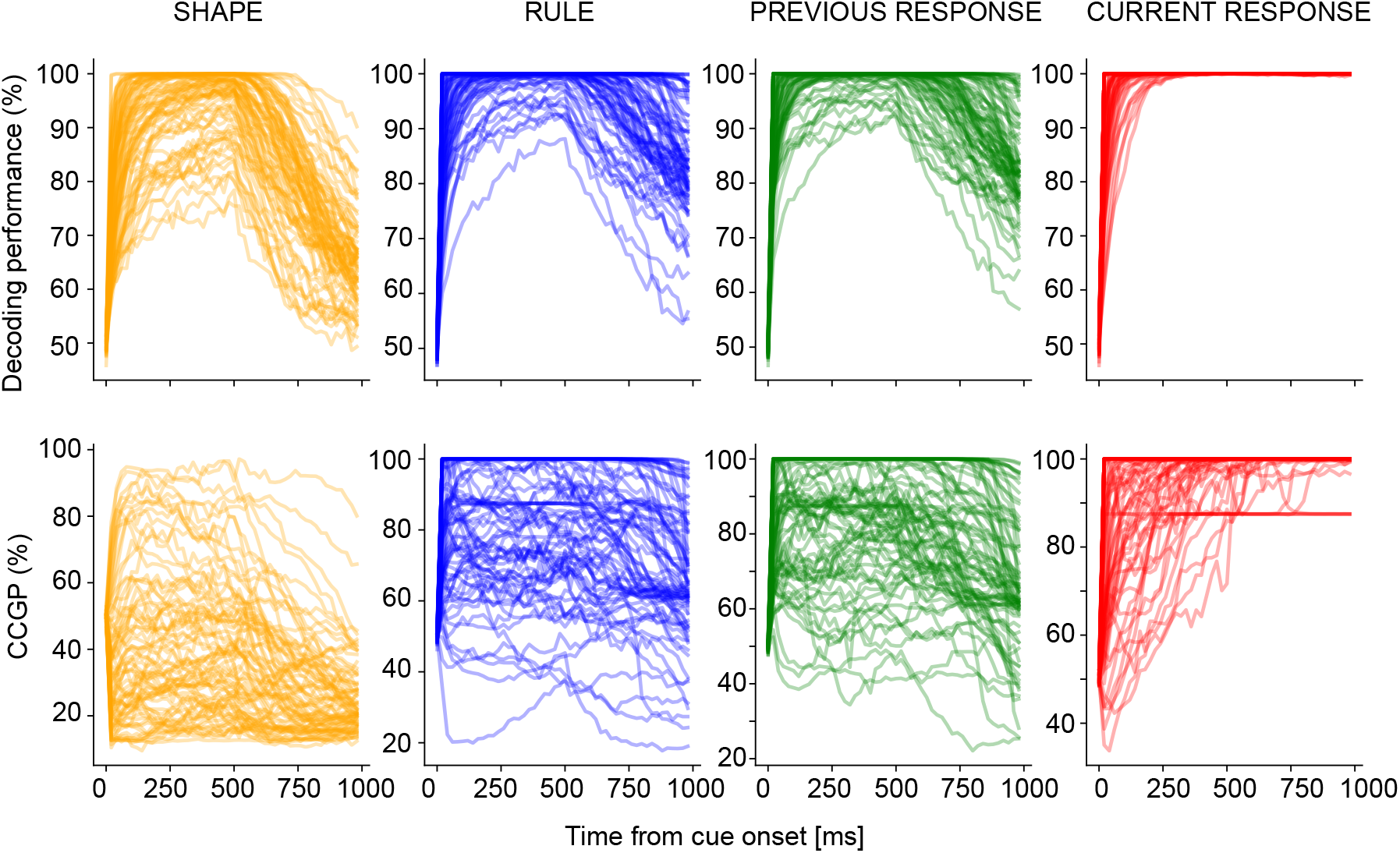
Decoding performance and CCGP of the four main task variables along time in all the trained RNNs. Top-row: Decoding performance for the shape (orange), rule (blue), previous (green), and current (red) response, from the cue onset. The cue offset is at 500*ms* from cue onset, with the first 500*ms* of the delay. Bottom-row: CCGP for the shape (orange), rule (blue), previous (green), and current (red) response, along time from the cue onset.

**Fig. 7:**
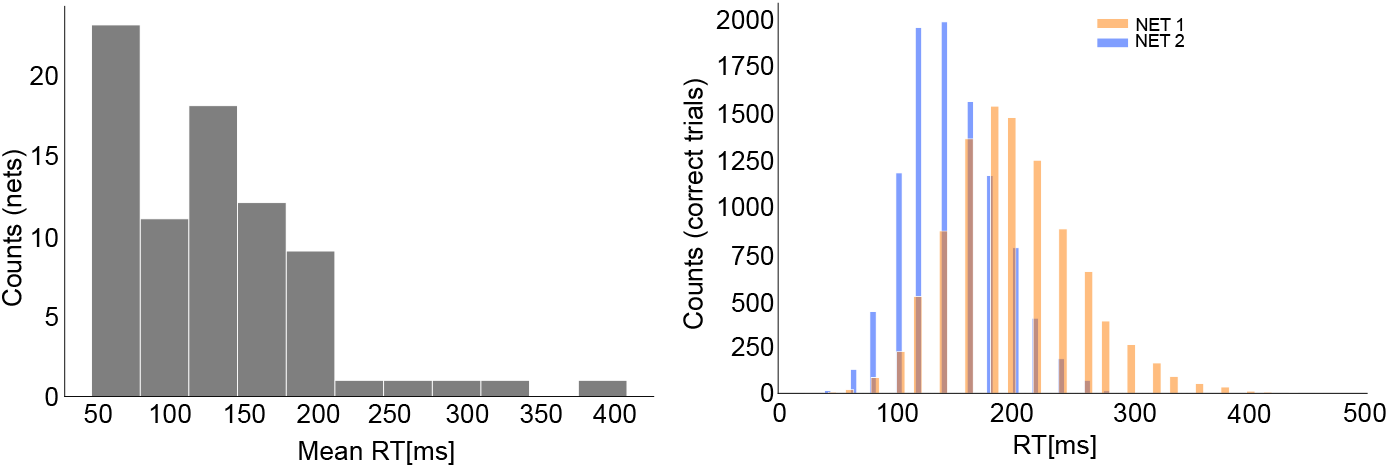
Distribution of reaction times across all the RNNs. Left: Distribution of mean reaction times (RTs) for all the trained recurrent neural networks (RNNs). Right: Distribution of RTs for NET 1 (orange) and NET 2 (blue) across the correct trials considered for the analyses. The mean of the distribution of RTs for NET 2 is smaller than the mean RTs in NET 1. This is due to the longer amount of training trials that allowed NET 2 to develop a better policy.

**Fig. 8:**
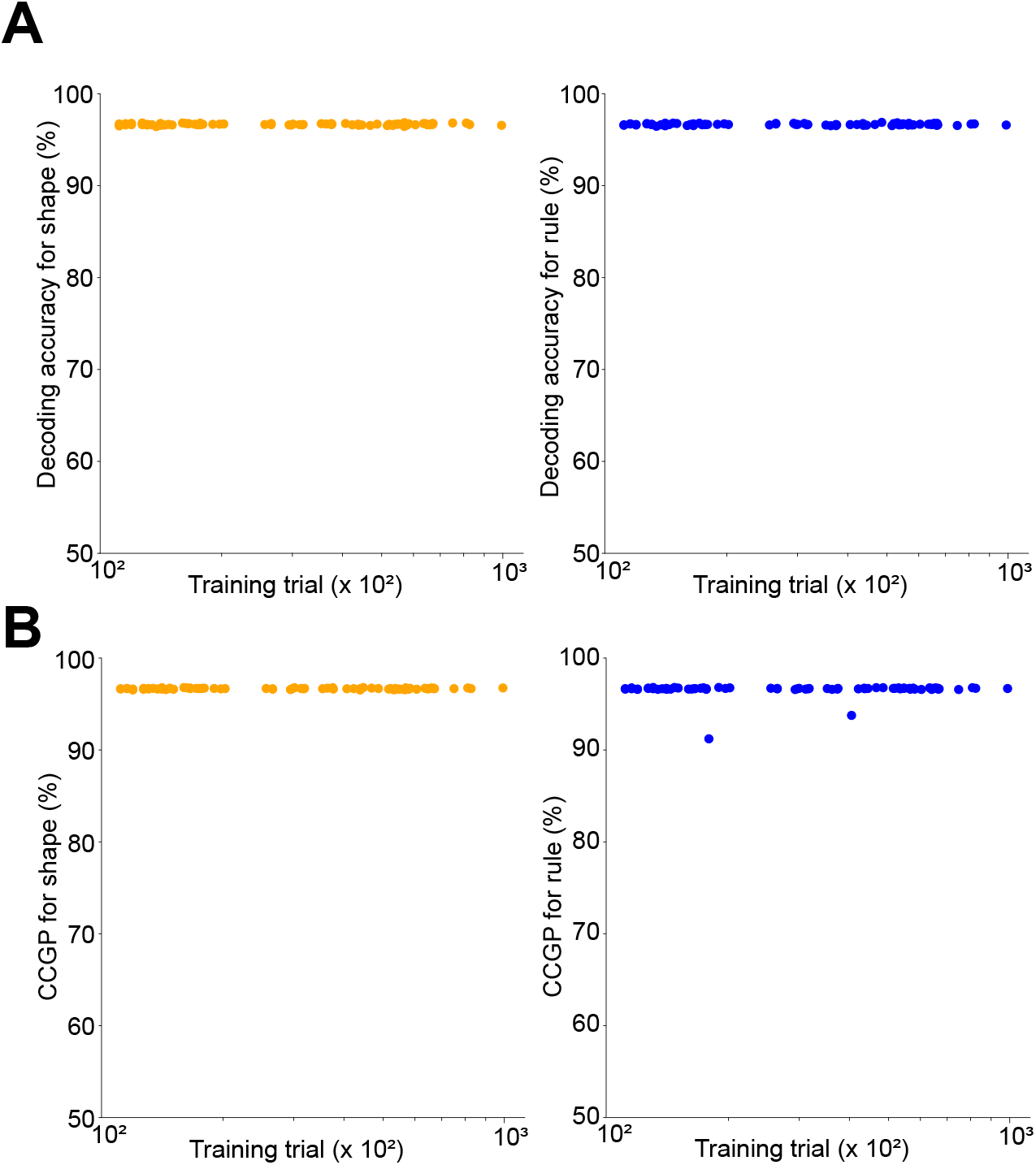
Decoding accuracy and CCGP for shape and rule in the input expansion layer of each RNN during the cue presentation as a function of the training trial. **A)** Decoding accuracy for shape (left) and rule (right) representation in the expansion input layer. Each point is a different RNN. The accuracy is very high for each of the RNNs without any significant difference. **B)** Same as in A but for the CCGP. In the expansion layer with random projection, the geometry still supports a high CCGP for both shape and rule across all the RNNs with no significant differences.

